# Hydrogen Bond Donors and Acceptors are Generally Depolarized in α-Helices as Revealed by a Molecular Tailoring Approach

**DOI:** 10.1101/536755

**Authors:** Hiroko X. Kondo, Ayumi Kusaka, Colin K. Kitakawa, Jinta Onari, Shusuke Yamanaka, Haruki Nakamura, Yu Takano

**Affiliations:** Kitami Institute of Technology, Kitami, Hokkaido 090-8507, Japan; Institute for Protein Research, Osaka University, Suita 565-0871, Japan; Graduate School of Science, Osaka University, Toyonaka 560-0043, Japan; Graduate School of Information Sciences, Hiroshima City University, Hiroshima 731-3194, Japan

## Abstract

Hydrogen-bond (H-bond) interaction energies in α-helices of short alanine peptides were systematically examined by precise DFT calculations, followed by a molecular tailoring approach (MTA). The contribution of each H-bond interaction in α-helices was estimated in detail from the entire conformation energies, and the results were compared with those in the minimal H-bond models, in which only H-bond donors and acceptors exist with the capping methyl groups. Consequently, the former interaction energies were always significantly weaker than the latter energies, when the same geometries of the H-bond donors and acceptors were applied. The chemical origin of this phenomenon was investigated by analyzing the differences among the electronic structures of the local peptide backbones of the α-helices and those of the minimal H-bond models. Consequently, we found that the reduced H-bond energy originated from the depolarizations of both the H-bond donor and acceptor groups, due to the repulsive interactions with the neighboring polar peptide groups in the α-helix backbone. The classical force-fields provide similar H-bond energies to those in the minimal H-bond models, which ignore the current depolarization effect, and thus they overestimate the actual H-bond energies in α-helices.

## Introduction

The hydrogen bond (H-bond) is one of the major factors that build the macromolecular structures of proteins, nucleic acids and their complexes. In particular, pair-wise H-bonds in protein backbones are essential to form their characteristic three-dimensional (3D) structures based on their ordered secondary structures, α-helices and β-sheets. Therefore, their structural energies should be correctly computed for analyses and predictions of protein 3D structures.

The individual force-fields used in classical molecular dynamics (MD) simulations show particular preferences and produce α-helical and β-structures^1–3)^. This phenomenon is usually not a problem for simulations of rigid globular protein structures, but it becomes a critical issue for folding simulations of flexible disordered regions^4,5)^ and long loops between secondary structures^6,7)^, to understand the functionally important conformational changes that occur as allosteric effects or induced folding upon ligand binding^4–7)^. Many attempts have been made to overcome this problem, by improving or rearranging the torsion energies^8–11)^, and by developing polarized charge models^12,13)^. However, these preferences have remained unclear, since their actual origins are unknown.

In our previous study^14)^, we computed the conformation energies of the secondary structures formed by peptide fragments, using several quantum chemical (QM) methods: the Hartree–Fock (HF) method, the second-order Møller-Plesset perturbation theory (MP2), and the density functional theory (DFT). Consequently, we found that a high quality DFT method including van der Waals interactions, B97D/6-31+G(d), was comparable to the MP2 method, which is reliable but time-consuming, for the Ace–(Ala)_*n*_–Nme system *in vacuo*^14)^. Using this DFT method, the energies of parallel and anti-parallel β-sheets can be approximated more or less by the classical force-fields, AMBER ff99SB^15)^, but those of the α-helical structures are significantly different. The differences were suggested to originate from the electrostatic energies associated with the H-bonds^14)^.

In this paper, by using the molecular tailoring approach (MTA)^16)^ with the DFT B97D/6-31+G(d) method, we dissected the individual interaction energy associated with each H-bond, to form typical α-helix backbones with different lengths. To analyze the origin of the H-bond interaction energy in an α-helix, we designed additional simplified models: a minimal H-bond (MH) model, composed of only the atoms forming a single H-bond, and a single-turn (ST) model, composed of three successive alanine residues, (Ala)_3_, in the α-helix capped by acetyl and N-methyl groups at the N- and C-termini, respectively. For those models, the individual H-bond energies were also computed by using MTA with the same DFT method, and the differences in the H-bond energies and the electronic structures among the complete α-helix and several models were analyzed. Finally, we discuss the putative reason underlying the secondary structure preferences in the classical force-fields used in molecular mechanical (MM) calculations.

## Materials and Methods

### α-helical structure (AH), Single turn (ST), and Minimal H-bond (MH) models

The α-helix models were first constructed by using 3- to 8-mer poly-alanine amino acids capped with an acetyl group (Ace) and an *N*-methyl amide group (Nme), denoted as Ace-(Ala)_*n*_-Nme, with the uniform (*φ, ψ, ω*) backbone angles for each residue: *φ* = −57°, *ψ* = −47°, and *ω* = 180°. Here, *n* is from 3 to 8. Each structure was optimized *in vacuo* by energy minimization of the electronic state, while maintaining the backbone angles as mentioned below.

There are one to (*n*−2) backbone hydrogen-bonds (H-bonds) in the optimized Ace-(Ala)_*n*_-Nme (3 ≤ *n* ≤ 8) α-helical structures, which are denoted here as “AH models”, between the backbone carbonyl group (-C=O) and the amide group (-NH) from the N- to C-terminus. They are denoted as AH*n*-1 to AH*n*-(*n*−2), respectively (Figure 1A), and their H-bond energies were individually computed by using MTA with the DFT method.

**Figure 1:**
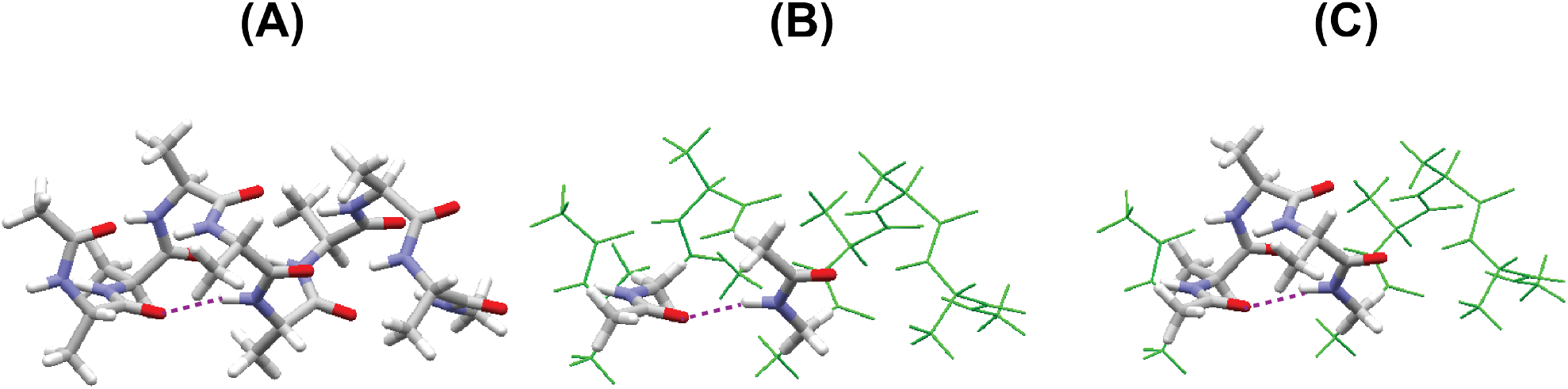
(A) The optimized α-helical structure for *n* = 8, α-helical structure (AH) model, (B) a minimal H-bond (MH) model, and (C) a single-turn (ST) model. In all of the models, the N-termini are capped by an acetyl group, Ace, and the C-termini are capped by an *N*-methyl group, Nme. Here, the target is the second H-bond, as indicated by the purple dotted lines.

To analyze the origin of the H-bond interaction energy in the α-helix, we designed a minimal H-bond model (MH model), which is composed of two separated *N*-methylacetamide molecules, mimicking a single H-bond between the *i*-th and (*i*+4)-th residues (Figure 1B). In addition, we designed a single-turn model (ST model), which is composed of three successive alanine residues, (Ala)_3_, in the α-helix capped by acetyl and *N*-methyl groups at the N- and C-termini, respectively (Figure 1C). The atom positions in these two models were the same as those in the corresponding AH models, except for the N- and C-terminal capping groups. For those models, the individual H-bond energies were computed by using MTA with the DFT method, in the same manner as for the AH models, and the energy differences among the H-bonds in those models were analyzed.

### Theoretical calculations

All calculations including the structure optimization were performed on the above atomic models with the Gaussian09 program packages^17)^ with the DFT B97D/6-31+G(d) method, which can correctly reflect the van der Waals interactions, and this method was confirmed to be comparable to the MP2 method for the Ace–(Ala)_*n*_–Nme system *in vacuo*^14)^.

It is not straightforward to extract the H-bond energy as a part of a large molecule, where the donor and acceptor atoms are linked through several covalent bonds. In fact, twelve covalent bonds link the acceptor atom, O, and the donor atom, H, in the backbone α-helical H-bond between the *i*-th and (*i*+4)-th amino-acid residues. Namely, the backbone *a*-helical H-bond is not a simple, additive pair-wise interaction, since it includes many-body effects with non-additive natures.

For that purpose, we employed the Molecular Tailoring Approach (MTA) developed by Deshmukh and Gadre, who showed that it is possible to estimate the intramolecular backbone H-bond energies of 3_10_-helices in several model polypeptides^16)^. Here, we used MTA to systematically compute the backbone H-bond energies in several α-helical peptide models.

The total energy of Ace-(Ala)_*n*_-NMe is approximated by dividing-and-conquering the energies of the fragments by MTA, using the following equation:

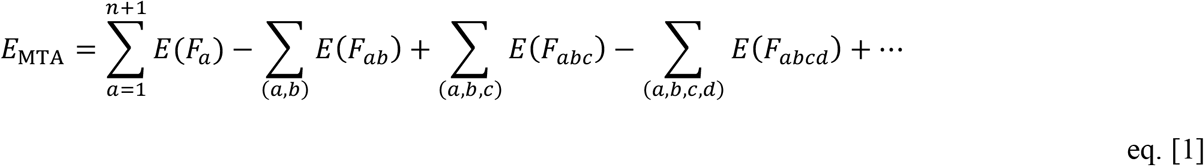

In eq. [1], each sum was taken for the possible combinations for (*a, b*, …) where 1 ≤ *a, b*, … ≤ (*n*+1). An example (*n* = 3) of the fragment models is shown in Figure S1. The total energies of the systems are well approximated by the combinations of all possible fragments, as described below, and thus we computed the energy of the entire system (*G*_0_ in Figure 2), instead of the combination in the original MTA method^16)^. We also computed the energy of a peptide fragment lacking the acceptor group (*G*_1_), the energy of a fragment lacking the donor group (*G*_2_), and the energy of a fragment lacking both the acceptor and donor groups (*G*_12_), as shown in Figure 2. The H-bond energy, *E*_HB_, and the electron density change upon H-bond formation, Δ*ρ*_MTA_, were estimated as follows:

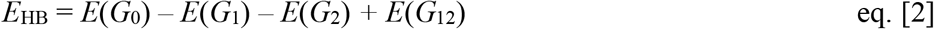

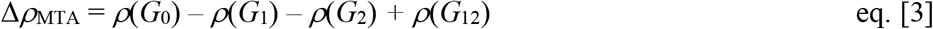

**Figure 2:**
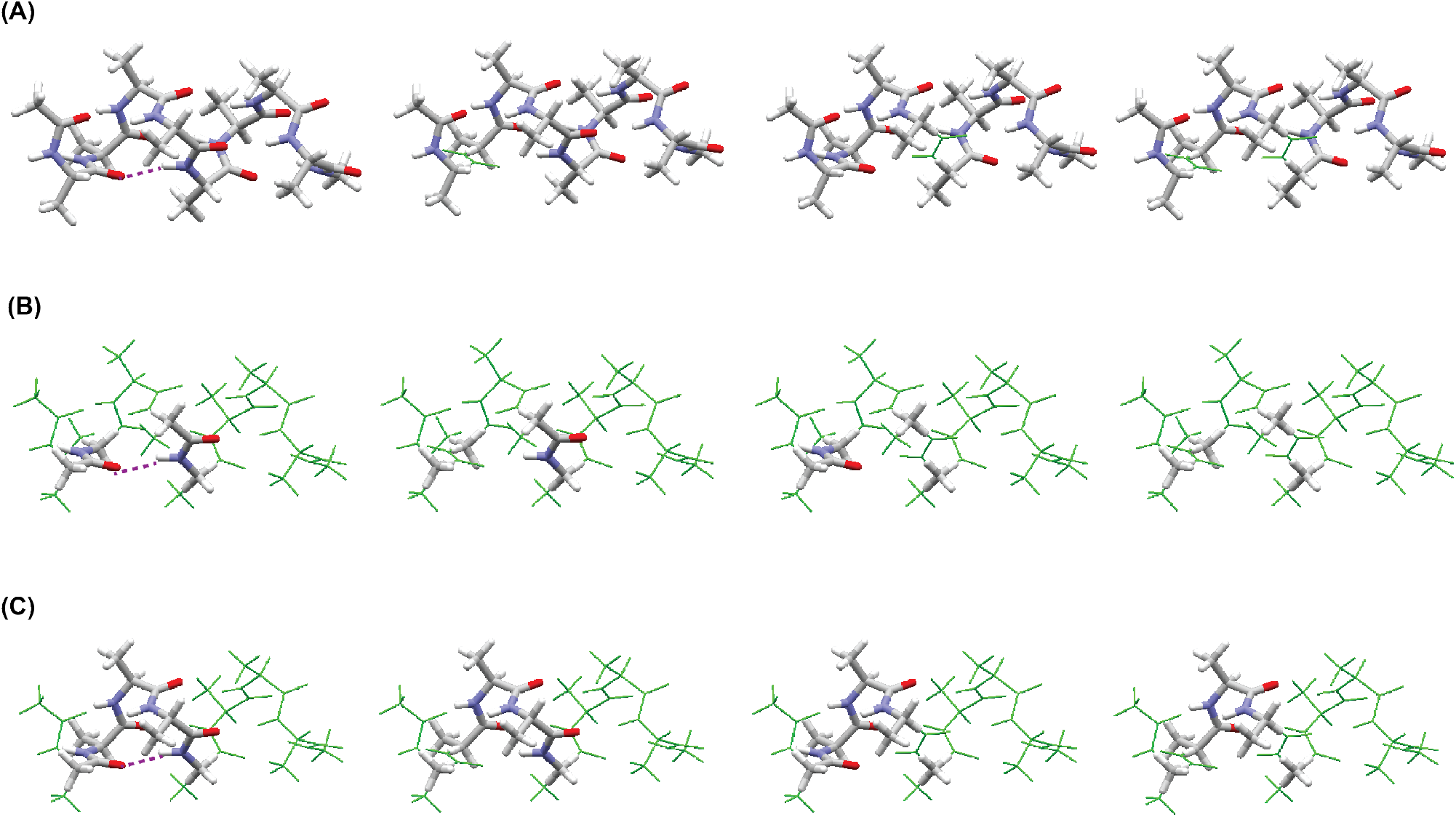
Flow of MTA computations for (A) the AH model, (B) the MH model, and (C) the ST model. The models are defined in Figure 1. In each picture, from the left to right, *G*_0_, *G*_1_, *G*_2_, and *G*_12_ are shown by sticks with CPK colors. The target here is the second H-bond, as indicated by the purple dotted lines. The thin green lines are the original Ace-(Ala)_8_-Nme structure.

We also calculated the stabilization energy (SE) in eq. [4], which we introduced in our previous paper^14)^, by using the same DFT B97D/6-31+G(d) method:

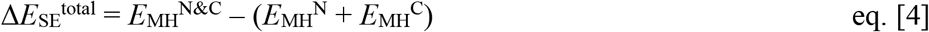

where *E*_MH_^N&C^ is the total energy of the MH model with both the N- and C-terminal *N*-methylacetamide molecules (Figure 1B). *E*_MH_^N^ and *E*_MH_^C^ are the total energies of the two separated *N*-methylacetamide molecules at the N- and C-termini, respectively. Namely, Δ*E*_SE_^total^ is the energy difference between the total energy of the MH model with the α-helical H-bond and that where the H-bond acceptor and the donor are separated infinitely. All of the SE values were corrected for the basis set superposition error (BSSE) by the counterpoise method of Boys and Bernardi^18)^. The ordinary electron density change upon H-bond formation, Δ*ρ*_SE_, was computed as follows:

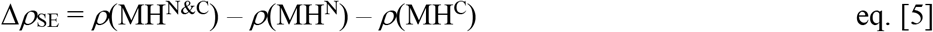

As references, we also computed the MM interaction energies for the corresponding H-bond interactions, using the following eq. [6]

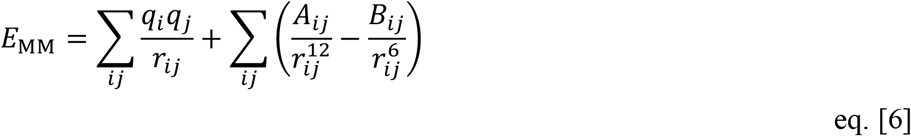

where *i* and *j* are four atoms attributed to peptides *I* and *J*, respectively: {C, O, N, and H}. *A_ij_* and *B_ij_* are the Lennard–Jones coefficients, *r_ij_* is the distance between *i*-th and *j*-th atoms, and *q_i_* is the atomic partial charge of the *i*-th atom. Here, AMBER ff99SB force-field parameters^15)^ were used.

Electron density changes were computed using the cube files in the Gaussian09 program packages^17)^, and the figures of the molecules with the electron density changes were produced by UCSF Chimera^19)^.

## Results

### H-bond energies in α-helices

The total energies of Ace-(Ala)_*n*_-Nme estimated by MTA (*E*_MTA_), computed by eq. [1], coincided well with the ordinary total energies *E*(*F*_0_) of the complete AH models. In fact, the differences in the values calculated by *E*(*F*_0_) and MTA, *E_MTA_* – *E*(*F*_0_), are indicated in Table 1, and all of them were less than 0.09 kcal/mol, representing about 0.00001% of the total energies. These differences are similar to that obtained in the previous study of the 3_10_-helix, where the difference was 0.11 kcal/mol^16)^.

**Table 1.** Total energies by the original DFT computation, *E*(*F*_0_), and the MTA method (*E*_MTA_) for the AH3, 4, 5, 6, 7, and 8 structures, corresponding to Ace-(Ala)_*n*_-Nme with *n* from 3 to 8. The differences are also shown.

| Structure | $E(F_0)$<br>(kcal/mol) | $E_{\text{MTA}}$<br>(kcal/mol) | $E_{\text{MTA}} - E(F_0)$<br>(kcal/mol) |
| --- | --- | --- | --- |
| AH3 | -621168.249 | -621168.157 | 0.092 |
| AH4 | -776275.937 | -776275.930 | 0.007 |
| AH5 | -931384.468 | -931384.465 | 0.003 |
| AH6 | -1086493.526 | -1086493.487 | 0.040 |
| AH7 | -1241602.973 | -1241602.883 | 0.090 |
| AH8 | -1396712.819 | -1396712.860 | -0.041 |

In Figure 3A, the H-bond energies between the backbone donors and acceptors are plotted for individual pairs for the models AH3 to AH8, depending on the distance between the donors and the acceptors for the AH models, ST models, and MH models by the MTA method, and the classical H-bond energies given by the MM calculation. The colors indicate the α-helical structures (Ace-(Ala)_*n*_-Nme) with different lengths (3 ≤ *n* ≤ 8), as indicated in the caption of Figure 3. The structural variations were caused by the energy minimization procedures for the entire α-helical conformations, as mentioned in the Methods section. The actual energy values are summarized in Table S1 in the supplementary material.

**Figure 3:**
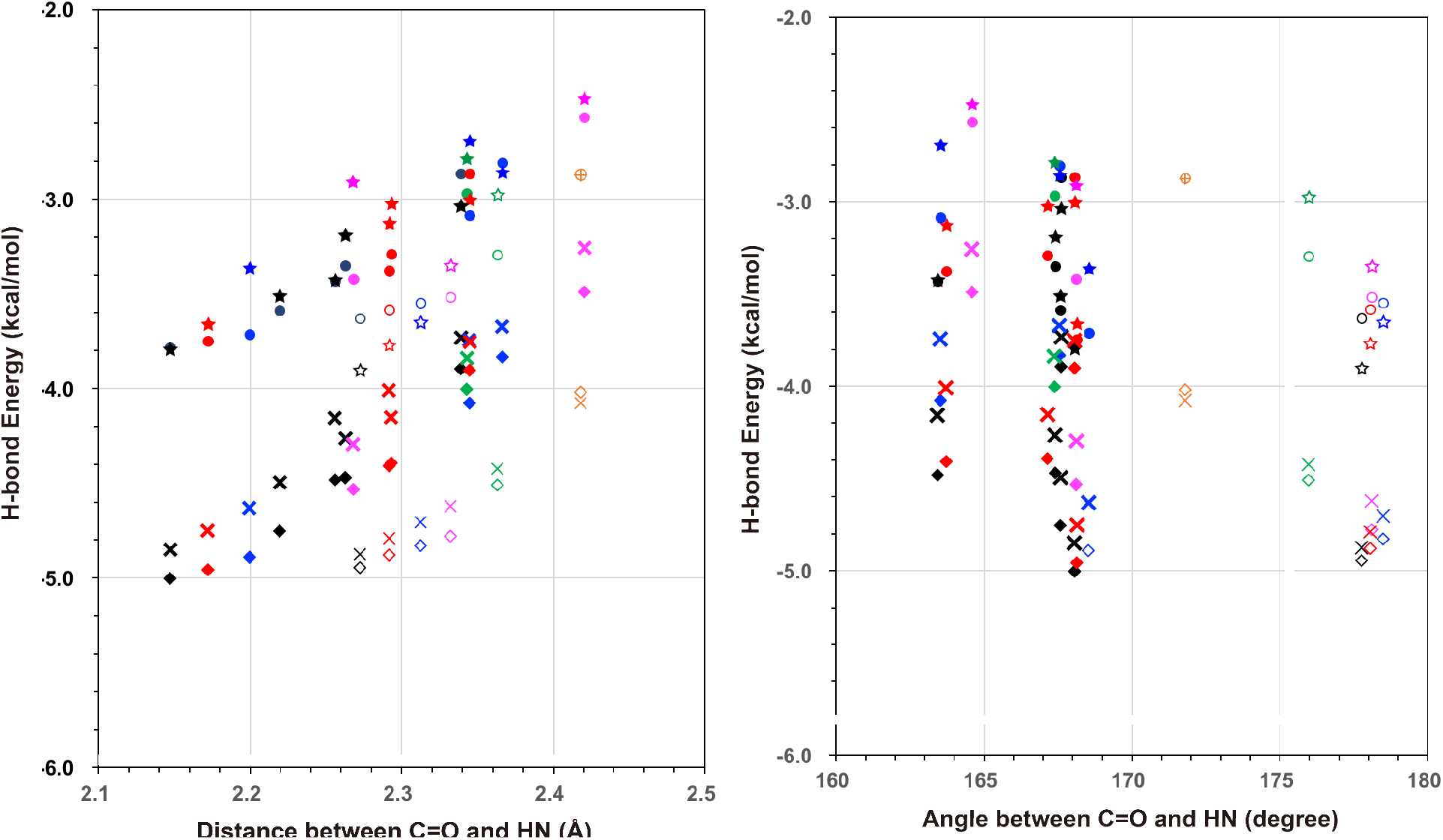
H-bond energy between the backbone donor and acceptor for each pair depending on (A) the distance between the O atom of the acceptor group, C=O, in the *i*-th residue, and the H atom of the donor group, NH, in the (*i*+4)-th residue, and on (B) the angle between the two vectors of C to O of the C=O and N to H of the NH. The resultant H-bond energies by the AH, ST, MT and MM models are indicated by the symbols ★, ●, ◆, and ×, respectively. The open symbols and thin marks are the values for the most N-terminal H-bond pairs. Others are shown by the filled symbols and thick marks. The symbol colors, orange, green, magenta, blue, red and black, are for the structures of AH3, AH4, AH5, AH6, AH7 and AH8, respectively.

As shown in Figure 3A and Table S1, the N-terminal helical turns were largely deformed from the initial conformations providing different H-bond lengths during the energy minimization procedure. Moreover, the angle between the two vectors of the carbon atom to the oxygen of the carbonyl group of the *i*-th residue and the nitrogen to the hydrogen of the amide group of the (*i*+4)-th residue was also deformed from the initial value at the N-terminus, as shown by the open and thin symbols in Figure 3B. Those vector pairs formed angles larger than 170°, and most of them were close to 180°. At the third H-bond, this angle was slightly smaller, by about 5°, than those of many other typical vector pairs, which were about 168° (Figure 3B).

Here, the H-bond energies in all of the models correlated well. In fact, when the H-bond energies calculated by the MTA method for the AH and ST models are plotted against those for the MH model, the H-bond energies calculated for the AH and ST models strongly correlated with that for the MH model, as shown in Figure 4, with Pearson’s correlation coefficients of 0.885 and 0.987, respectively.

**Figure 4:**
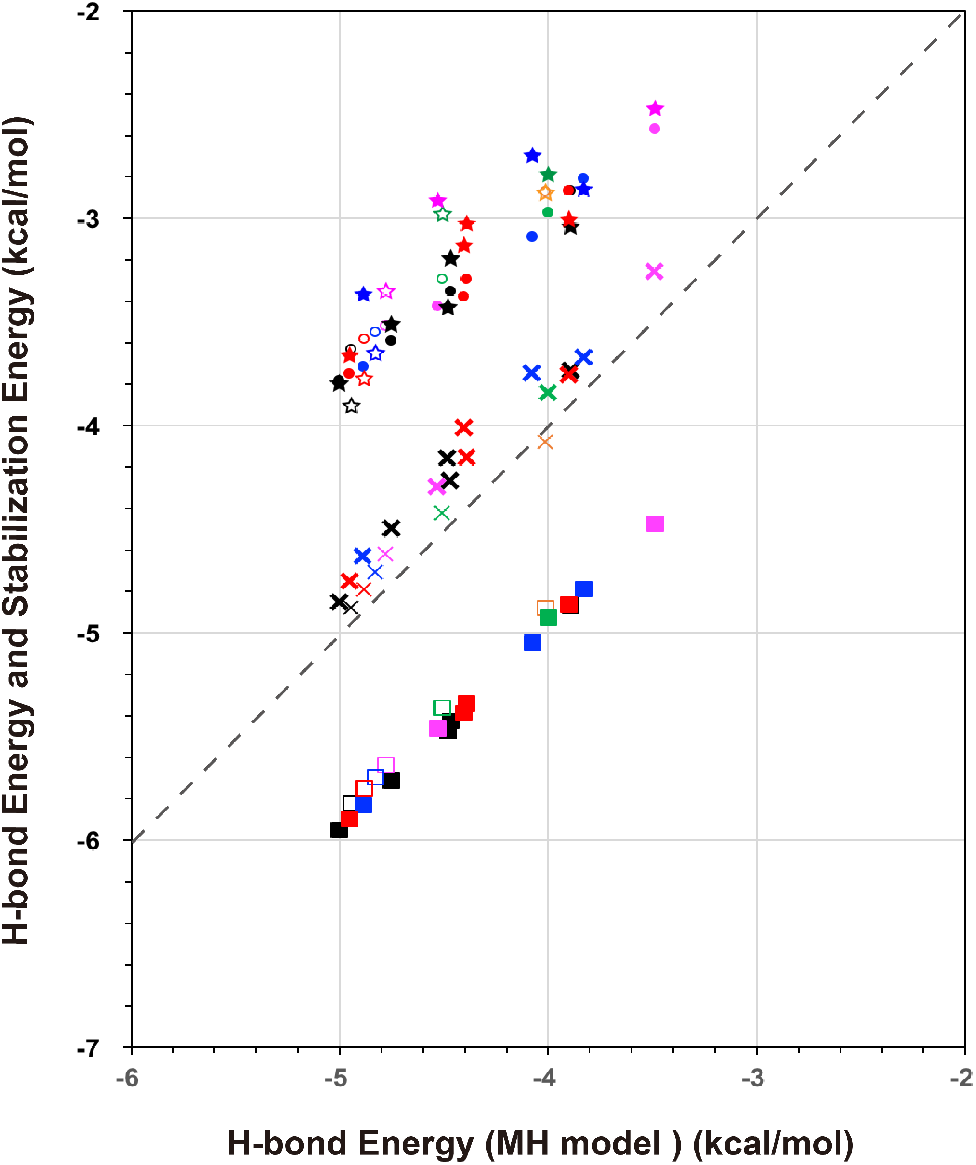
Correlations of the H-bond energies in the AH model (★), the ST model (●), the MM model (×), and the stabilization energy (SEs: ■) against those by the MH model. The meanings of the filled and open symbols with different colors are the same as those in the caption of Figure 3. The dashed line shows a guide where the longitudinal axis values have the same H-bond energies by the MH models.

The MM values have also a high correlation coefficient, 0.975, with the MH model, and even the absolute MM values are very close to the H-bond energies calculated by the MH models. In contrast, the H-bond energies obtained by both the AH and ST models remarkably deviated from those calculated by the MH models.

### Stabilization energies (SEs) in MH models

The SE values defined in eq. [4] for the MH models, Δ*E*_SE_^total^, are also plotted in Figure 4, and the actual values are provided in Table S1. The SE values correlated very well with the corresponding H-bond energies in the MH models, with a Pearson’s correlation coefficient of 0.995. The SE values were 0.93 kcal/mol lower than the corresponding H-bond energies in the MH models except for the N-termini, where the N-terminal backbone structures were largely deformed and their SE values were 0.86 kcal/mol lower than the H-bond energies in the MH models.

### Electronic structures around the H-bond donors and acceptors

In addition to the H-bond energies, the MTA method can approximate the electronic structures around the H-bond donors and acceptors in α-helices. In fact, the ordinary electron density change upon H-bond formation, Δ*ρ*_SE_, for the first H-bonding donor and acceptor groups (8-1) of the α-helical alanine octamer, Ace-(Ala)_8_-Nme, was provided by eq. [5], and it is shown in Figure 5 (A). The corresponding electron density changes for the AH, MH, and ST models, Δ*ρ*_MTA_, computed by eq. [3], are also shown in Figure 5 (B)-(D), respectively. It is obvious that the Δ*ρ*_MTA_ values are all similar to the Δ*ρ*_SE_ values, where the electron density increases around the oxygen atom of the carbonyl (C=O) group at the *i*-th residue and decreases around the hydrogen atom of the amide (N-H) group at the (*i*+4)-th residue. For the other structures from (8-2) to (8-6), the Δ*ρ*_MTA_ values of the AH models are shown in Figure S2 (A)-(E).

**Figure 5:**
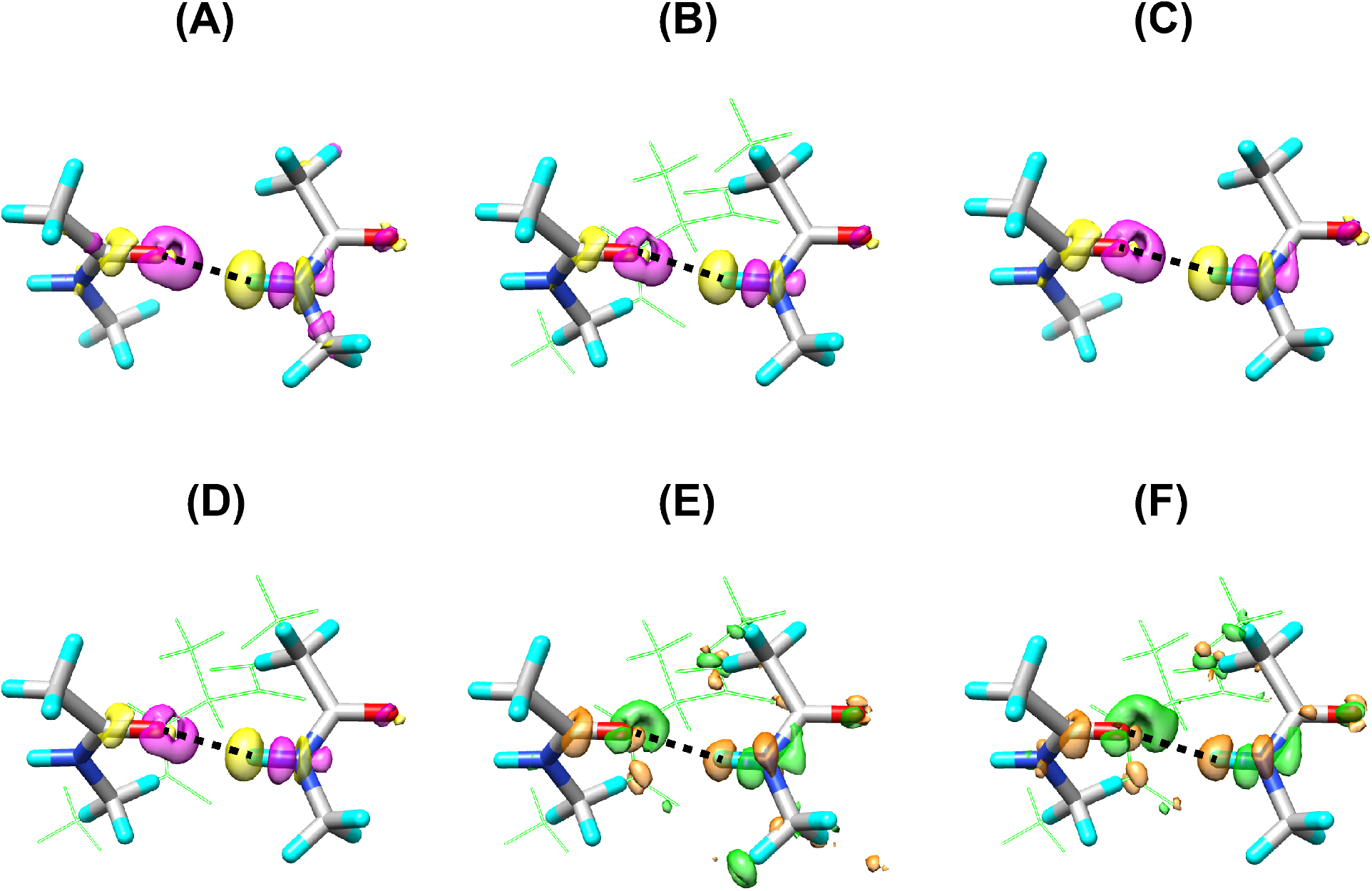
(A) The electron density change upon H-bond formation, Δ*ρ*_SE_, for the α-helix AH8-1 structure provided by eq. [5]. The corresponding electron density changes, Δ*ρ*_MTA_, computed by eq. [3] for the (B) AH, (C) MH, and (D) ST models. The yellow surface is the contour surface at −0.001 au, and the magenta one is that at 0.001 au. (E) The difference in the electron density change between the AH and MH models ΔΔ*ρ*_MTA_^AH−MH^ by eq. [7], and (F) that between the ST and MH models ΔΔ*ρ*_MTA_^ST−MH^ by eq. [8]. The green surface is the contour surface at −0.0002 au, and the orange one is that at 0.0002 au. The atoms in the MH model are shown by stick models, and the other atoms in the AH and ST models are shown by open green sticks. The black dotted line is the H-bond between the oxygen atom of the C=O group at the *i*-th residue and the hydrogen atom of the N-H group at the (*i*+4)-th residue.

The differences in the electron density changes between the AH and MH models, and those between the ST and MH models were further computed, respectively:

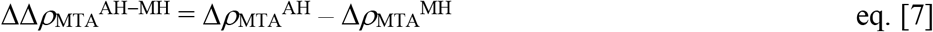

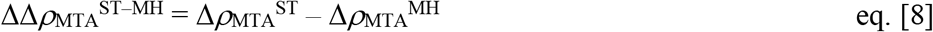

In Figure 5(E) and (F), the ΔΔ*ρ*_MTA_^AH−MH^ and ΔΔ*ρ*_MTA_^ST−MH^ values for the first α-helical turn (8-1) are shown between the AH and MH models and between the ST and MH models, respectively. The electron density near the oxygen atom of the C=O group at the *i*-th residue decreased in both the AH and ST models, as compared with that in the MH model. In contrast, the electron density near the hydrogen atom of the N-H group at the (*i*+4)-th residue increased in the AH and ST models, as compared with that in the MH model. The differences in the electron density changes upon H-bond formation for the AH and ST models, ΔΔ*ρ*_MTA_^AH−MH^ and ΔΔ*ρ*_MTA_^ST−MH^, in all structures from 8-2 to 8-6 are shown in Figures S3 and S4, respectively.

## Discussion

### H-bond energies by the MTA method

The energies of Ace-(Ala)_*n*_-Nme provided by the MTA method are approximated values. However, as shown in Table 1, the differences in the values between the ordinary total energies for the complete *F*_0_ and those obtained by MTA, *E*_MTA_ – *E*(*F*_0_), are very small. They are also small even in comparison to the H-bond energies, which are about −3 to 4 kcal/mol in this study. Thus, the H-bond energies estimated by the current MTA method should be quantitatively reliable, with about 3% errors, for discussing the H-bond interactions in α-helices.

The accuracy of the MTA method largely relies on the borders separating molecular segments. Historically, this issue was recognized as the “nearsightedness of electronic matter (NEM)”^20)^ to divide-and-conquer large molecular systems in general. Using the theoretical computations on the basis of the linear response function, the sp^3^ junction was the most suitable location for partitioning peptide systems^21,22)^. In the current MTA procedures, all of the fragmentations followed the sp^3^ junction mechanism to block the propagation of the electron density deviation.

In the MH models where the two peptide groups for hydrogen donors and acceptors are separated without any covalent bonds, the H-bond energies should directly correspond to the Stabilization Energies (SEs) including the BSSE corrections^18)^. In fact, an almost perfect correlation appeared between the H-bond energies and the SEs, as shown in Figure 4 and Table S1, and the differences were always the same, 0.93 kcal/mol, except for the N-termini where the backbone structures were largely deformed during the energy optimization procedures. Those differences are due to other interactions among the methyl groups that capped the N- and C-terminal peptide groups than the H-bond energies given by the current MTA analysis using the MH models. In addition, as shown in Figure 5 (A) and (C), the change in the electron density upon H-bond formation, Δ*ρ*_SE_ in eq. [5], is also well approximated by DΔ*ρ*_MTA_^MH^ in eq. [3].

Thus, the current MTA method provides good approximations of the H-bond energies and electronic structures in MH models, and so it is expected to give a reliable analysis for the H-bond interactions of the AH and ST models as well, as shown in Figure 5 (B) and (D). Morozov *et al*.^23)^ reported a similar approach to analyze the cooperativity of α-helix formation by using separated α-helical peptide fragments. In particular, their model including the short-range contribution was, in principle, designed to compute the dimerization energies using the SEs. Namely, their short-range interactions did not correctly account for the non-additive many-body interactions, and ignored the effects of the α-helical backbone atoms linking the H-bond acceptor and donor.

### Analysis of the electronic structures

It is clear from Figure 4 and Table S1 that the H-bond energies for the AH models have similar values to those for the ST models, although the former ones tend to be slightly weaker than the latter ones. In contrast, the H-bond energies in the AH and ST models significantly deviated from those in the MH models, although the Pearson’s correlation coefficients were very high. As shown in Figures 5E, 5F, S3 and S4, the electronic structures around the H-bonds in the AH and ST models distinctively deviated from those in the MH models, with the depolarization of the hydrogen donor and acceptor groups.

Thus, the phenomenon should be caused by the precise electronic structures in the ST and AH models. In order to analyze the origin of this phenomenon, we focused on the six model structures of AH8-1 to AH8-6. We found that the distances between the oxygen atoms in the carbonyl group of the *i*-th and (*i*+1)-th residues in the six H-bond pairs are short, 3.510±0.144 Å. In addition, those between the hydrogen atoms in the amide group of the (*i*+3)-th and (*i*+4)-th residues are also short, 2.676±0.038 Å. These short distances suggest that the carbonyl oxygen of the *i*-th residue has less electron density, and the amide hydrogen of (*i*+4)-th residue has more electron density, as revealed in Figure 5E and 5F.

By applying the Hirshfeld population analysis^24–26)^, the electronic structures in the six ST models were analyzed around the carbonyl oxygen of the *i*-th residue and the amide hydrogen of the (*i*+4)-th residue, in comparison to those in the MH models, in which no neighboring carbonyl (C=O) or amide (NH) groups exist. As shown in Table S2C, for the *G*_2_ fragment of the ST model (Figure 2C), which lacked the H-bond donor group at the (*i*+4)-th residue but included the effect of the C=O group of the (*i*+1)-th residue, the Hirshfeld atomic charge of the carbonyl oxygen of the *i*-th residue was 0.0303e±0.0036e larger than that in the MH model. The contribution of the amide hydrogen of the (*i*+4)-th residue for the *G*_1_ fragment of the ST model, which lacked the H-bond acceptor group at the *i*-th residue but included the effect of the amide group of the (*i*+3)-th residue, was 0.0154e±0.0010e less than that in the MH model. Similar Hirshfeld charge changes were also observed for the entire *G*_0_ systems, where H-bonds are formed between the C=O group of the *i*-th residue and the NH group of the (*i*+4)-th residue, as shown in Table S2A.

These depolarization effects also correlate with the local dipole moments. The local interatomic dipole moment for a system composed of *N* charges {*q_k_* | *k* = 1, …, *N*} is described by eq. [9]

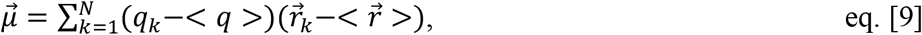

where <*q* > is the average of the Hirshfeld atomic charges and 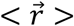 is their center position. For the C=O group of the *i*-th residue and the NH group of the (*i*+4)-th residue, the local dipole moments become simple, as shown in eqs. [10] and [11]:

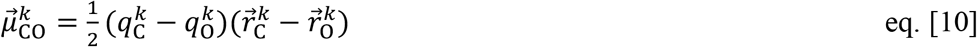

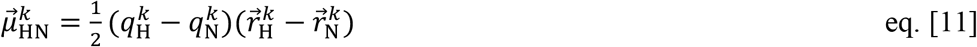

Here, *q^k^*_C_ and *q^k^*_O_ are the Hirshfeld atomic charges of the C and O atoms in the C=O group of the *k*-th residue, and *q^k^*_H_ and *q^k^*_N_ are those of the NH group of the *k*-th residue, respectively. 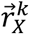 is the position vector of the corresponding atom *X* of the *k*-th residue. Hereafter, only the absolute values, 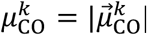 and 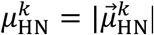, are used for the following discussion.

Due to the closely located electric dipole *μ*^*i*+1^_CO_ at the (*i*+1)-th residue to *μ^i^*_CO_ at the *i*-th residue in the parallel direction, when an α-helical conformation is formed, the dipole moments should decrease by their repulsive interaction. In the same way, due to the closely located electric dipole *μ*^*i*+3^_HN_ at the (*i*+3)-th residue to *μ*^*i*+4^_HN_ at the (*i*+4)-th residue in the parallel direction in an α-helix, those dipole moments should also decrease. From the Hirshfeld population analysis, as shown in Table 2B, the average ratio of *μ^i^*_CO_ in the ST model to that in the MH model was 0.938±0.002, and the average ratio of *μ*^*i*+4^_HN_ in the ST model to that in the MH model was 0.917 ±0.002, when the H-bonds were not formed between the C=O group of the *i*-th residue and the NH group of the (*i*+4)-th residue. As shown in Table 2A, when H-bonds were formed between the C=O group of the *i*-th residue and the NH group of the (*i*+4)-th residue, the average ratio of *μ^i^*_CO_ in the ST model to that in the MH model was 0.944±0.002, and the average ratio of *μ*^*i*+4^_HN_ in the ST model to that in the MH model was 0.946±0.003. Namely, these dipole moments were reduced by about 5–6 and 5–8%, respectively, depending on the backbone α-helical conformation. Although such population analyses may include some ambiguities for the absolute values of the atomic charges and dipole moments, these tendencies accurately reflect the changes of the electronic structures shown in Figures 5F and S4, as the depolarization effect.

**Table 2.** Inter-atomic dipole moments of the carbonyl group (C=O) of the *i*-th residue and those of the amide group (NH) of the (*i*+4)-th residue by the Hirshfeld population analysis. (A) ST, AP_N_, AP_C_, and MH models in *G*_0_ fragments. (B) ST, AP_N_, and MH models in *G*_2_ fragments, and ST, AP_C_, and MH models in *G*_1_ fragments.

| Dipole moments involved in H-bond |  | Dipole moments (Debye) for structures |  |  |  |  |  | Average (Debye) |
| --- | --- | --- | --- | --- | --- | --- | --- | --- |
|  |  | 8-1 | 8-2 | 8-3 | 8-4 | 8-5 | 8-6 |  |
| $\mu_{\text{CO}}^{i\text{-th}}$ of $i$-th residue in $G_0$ fragment | ST model | 1.3315 | 1.3055 | 1.3175 | 1.3187 | 1.3205 | 1.3277 | 1.3202 |
|  | MH model | 1.4163 | 1.3838 | 1.3913 | 1.3960 | 1.3994 | 1.4043 | 1.3985 |
|  | Ratio (ST/MH)<sup>1)</sup> | 0.9401 | 0.9434 | 0.9470 | 0.9446 | 0.9436 | 0.9455 | 0.9440 |
|  | AP<sub>N</sub> model | — | 1.3569 | 1.3566 | 1.3512 | 1.3594 | 1.3653 | 1.3579 |
|  | Ratio (AP<sub>N</sub>/MH)<sup>2)</sup> | — | 0.9806 | 0.9751 | 0.9679 | 0.9714 | 0.9722 | 0.9734 |
| $\mu_{\text{HN}}^{i+4\text{-th}}$ of $(i+4)$-th residue in $G_0$ fragment | ST model | 0.5477 | 0.5356 | 0.5375 | 0.5376 | 0.5426 | 0.5412 | 0.5404 |
|  | MH model | 0.5778 | 0.5642 | 0.5704 | 0.5691 | 0.5729 | 0.5740 | 0.5714 |
|  | Ratio (ST/MH)<sup>3)</sup> | 0.9479 | 0.9493 | 0.9423 | 0.9446 | 0.9471 | 0.9429 | 0.9457 |
|  | AP<sub>C</sub> model | 0.5691 | 0.5570 | 0.5623 | 0.5611 | 0.5632 | — | 0.5625 |
|  | Ratio (AP<sub>C</sub>/MH)<sup>4)</sup> | 0.9849 | 0.9872 | 0.9858 | 0.9859 | 0.9831 | — | 0.9854 |

1) Ratio of the absolute value of the dipole moment $\mu_{\text{CO}}^i$ of the $i$ -th residue in the ST model versus that in the MH model.
2) Ratio of the absolute value of the dipole moment $\mu_{\text{CO}}^i$ of the $i$ -th residue in the AP<sub>N</sub> model versus that in the MH model.
3) Ratio of the absolute value of the dipole moment $\mu_{\text{HN}}^{i+4}$ of the $(i+4)$ -th residue in the ST model versus that in the MH model.
4) Ratio of the absolute value of the dipole moment $\mu_{\text{HN}}^{i+4}$ of the $(i+4)$ -th residue in the AP<sub>C</sub> model versus that in the MH model.

(B) ST, AP<sub>N</sub>, and MH models in $G_2$ fragments, and ST, AP<sub>C</sub>, and MH models in $G_1$ fragments.
| Dipole moments involved in H-bond |  | Dipole moments (Debye) for structures |  |  |  |  |  | Average (Debye) |
| --- | --- | --- | --- | --- | --- | --- | --- | --- |
|  |  | 8-1 | 8-2 | 8-3 | 8-4 | 8-5 | 8-6 |  |
| $\mu_{\text{CO}}^i$ of $i$-th residue in $G_2$ fragment | ST model | 1.3404 | 1.3274 | 1.3324 | 1.3349 | 1.3352 | 1.3413 | 1.3353 |
|  | MH model | 1.4346 | 1.4166 | 1.4157 | 1.4224 | 1.4235 | 1.4256 | 1.4231 |
|  | Ratio (ST/MH)<sup>5)</sup> | 0.9343 | 0.9370 | 0.9412 | 0.9385 | 0.9380 | 0.9409 | 0.9383 |
|  | AP<sub>N</sub> model | – | 1.3856 | 1.3765 | 1.3729 | 1.3791 | 1.3824 | 1.3793 |
|  | Ratio (AP<sub>N</sub>/MH)<sup>6)</sup> | – | 0.9781 | 0.9723 | 0.9652 | 0.9688 | 0.9697 | 0.9708 |
| $\mu_{\text{HN}}^{i+4}$ of $(i+4)$-th residue in $G_1$ fragment | ST model | 0.5801 | 0.5826 | 0.5796 | 0.5797 | 0.5827 | 0.5781 | 0.5805 |
|  | MH model | 0.6333 | 0.6333 | 0.6326 | 0.6329 | 0.6339 | 0.6304 | 0.6327 |
|  | Ratio (ST/MH)<sup>7)</sup> | 0.9160 | 0.9199 | 0.9162 | 0.9159 | 0.9192 | 0.9170 | 0.9174 |
|  | AP<sub>C</sub> model | 0.6189 | 0.6190 | 0.6179 | 0.6180 | 0.6178 | – | 0.6183 |
|  | Ratio (AP<sub>C</sub>/MH)<sup>8)</sup> | 0.9773 | 0.9774 | 0.9768 | 0.9765 | 0.9746 | – | 0.9765 |

5) Ratio of the absolute value of the dipole moment $\mu_{\text{CO}}^i$ of the $i$ -th residue in the ST model versus that in the MH model.
6) Ratio of the absolute value of the dipole moment $\mu_{\text{CO}}^i$ of the $i$ -th residue in the AP<sub>N</sub> model versus that in the MH model.
7) Ratio of the absolute value of the dipole moment $\mu_{\text{HN}}^{i+4}$ of the $(i+4)$ -th residue in the ST model versus that in the MH model.
8) Ratio of the absolute value of the dipole moment $\mu_{\text{HN}}^{i+4}$ of the $(i+4)$ -th residue in the AP<sub>C</sub> model versus that in the MH model.

In order to examine the above phenomena, two other models, the HT_N_ (N-terminal Half-Turn) and HT_C_ (C-terminal Half-Turn) models, were constructed. In the HT_N_ model, the (*i*+2)-th and (*i*+3)-th Ala residues were both deleted from the ST model and capped by methyl groups, as shown in Figure S5 (A). In the HT_C_ model, the (*i*+1)-th and (*i*+2)-th Ala residues were both deleted, as shown in Figure S5 (B). Namely, the carbonyl group of the (*i*+1)-th residue is included in the HT_N_ model, and the amide group of the (*i*+3)-th residue is included in the HT_C_ model. Consequently, the H-bond energies for the HT_N_ and HT_C_ models were located in the middle between the corresponding ST and MH models, as shown in Figure 6 by the filled and open triangles, respectively. The actual energy values are summarized in Table S3.

**Figure 6:**
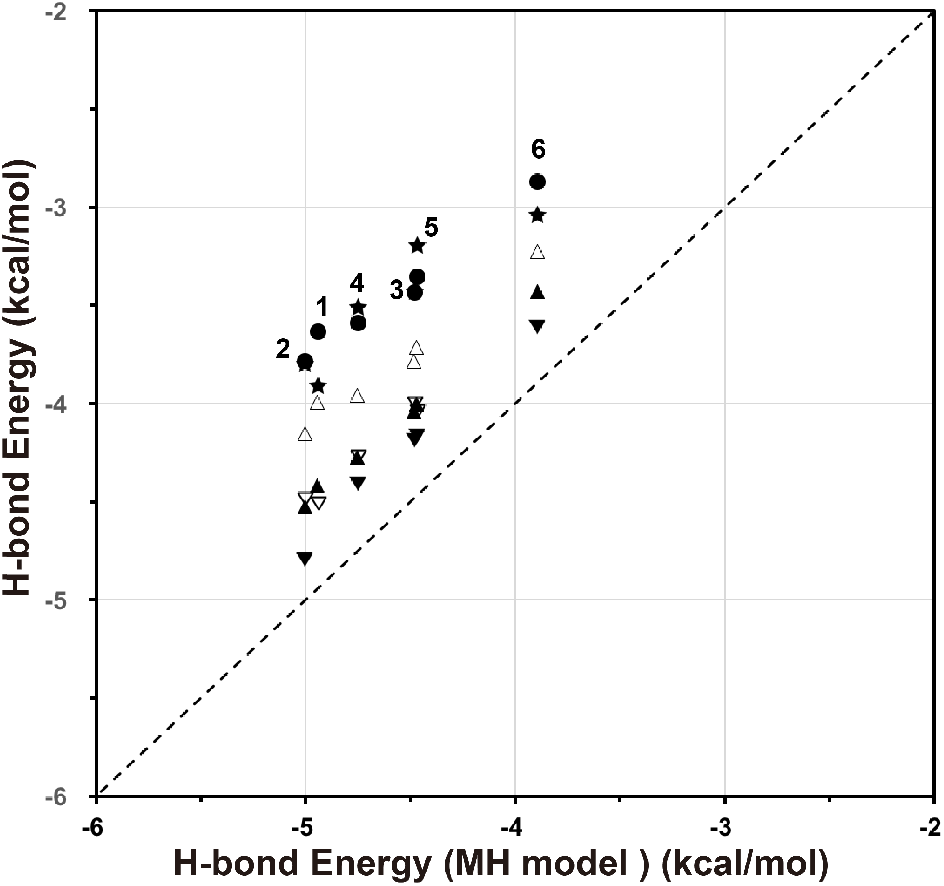
H-bond energies in the HT_N_ (N-terminal Half-Turn) model (▲), the HT_C_ (C-terminal Half-Turn) model (Δ) with those in the ST model (●), AH model (★) for the 8-1 to 8-6 structures, H-bond energies in AP_N_ (N-terminal additional peptide) model (▼) for the 8-2 to 8-6 structures, and those in the AP_C_ (N-terminal additional peptide) model (▽) for the 8-1 to 8-5 structures. The numbers show the location, *i*, of the H-bond from 1 to 6. The dashed line is a diagonal guideline.

A careful investigation of the H-bond energies of the AH and ST models in Table S1 and Figure 6 revealed that the H-bond energies in the AH models are always slightly weaker than those in the ST models, except for the N-termini or C-termini. These effects can be caused by the successive carbonyl group of the (*i*−1)-th residue and the amide group of the (*i*+5)-th residue, in a similar manner to the effects of the successive carbonyl group of the (*i*+1)-th residue and the amide group of the (*i*+3)-th residue in the opposite directions. In fact, the distances between the oxygen atoms in the C=O groups of the (*i*−1)-th and *i*-th residues in the five H-bond pairs are 3.506±0.037 (Å), and those between the hydrogen atoms in the NH groups of the (*i*+4)-th and (*i*+5)-th residues are 2.714±0.038 (Å), as shown in Table S2 (B).

Two additional models, the AP_N_ (N-terminal Additional Peptide) and AP_C_ (C-terminal Additional Peptide) models, were also considered, where an Ace-Ala group and an Alα-Nme group were added to the N-terminus and C-terminus of the MH model, respectively (Figure S5 (C) and (D)). The AP_N_ model has an interaction between the successive C=O groups at the (*i*-1)-th and *i*-th residues, and the AP_C_ model has another interaction between the successive NH groups at the (*i*+4)-th and (*i*+5)-th residues. Their H-bond energies computed by the MTA method are shown in Figure 6 and Table S3. Both of them have the middle H-bond energies between the corresponding ST and MH models, suggesting another depolarization effect.

These phenomena were also analyzed with the local electric dipole *μ^i^*_CO_ at the *i*-th residue in the AP_N_ model, and *μ*^*i*+4^_HN_ at the (*i*+4)-th residue in the AP_C_ model. Consequently, the ratio of *μ^i^*_CO_ in the AP_N_ model to that in the MH model was 0.971±0.004, and the ratio of *μ*^*i*+4^_HN_ in the AP_C_ model to that in the MH model was 0.977±0.001, when the H-bonds were not formed between the C=O group of the *i*-th residue and the NH group of the (*i*+4)-th residue (Table 2B). When H-bonds were formed between the C=O group of the *i*-th residue and the NH group of the (*i*+4)-th residue, the ratio of *μ^i^*_CO_ in the AP_N_ model to that in the MH model was 0.973 ±0.004, and the ratio of *μ*^*i*+4^_HN_ in the AP_C_ model to that in the MH model was 0.985±0.001 (Table 2A). These dipole moments were both also reduced by about 2−3%, and their contributions to the total H-bond energies are smaller than those of *μ*^*i*+1^_CO_ or *μ*^*i*+3^_HN_.

As shown in Figure 6 and Table S3, the H-bond energies due to the surrounding carbonyl groups around the *i*-th residue and those of the amide group around the (*i*+4)-th residue were not additive. Namely, the simple summations of the energy differences of the H-bond energies between the HT_N_ and MH models and between the HT_C_ and MH models are always larger than the direct differences between the ST and MH models. Similarly, simple summations of the differences in the H-bond energies between the AP_N_ and MH models and between the AP_C_ and MH models could also overestimate the differences between the AH and ST models. Thus, they should be considered as non-additive many-body effects.

Finally, the putative effects of the helical dipoles at the backbone peptide planes that are far from the target H-bond locations were also investigated. Although the neighboring peptide dipoles at the (*i*−1)-th and (*i*+5)-th residues slightly contributed to the H-bond energy of the unit α-helix from the *i*-th to (*i*+4)-th residues, as discussed above, there was no significant dependence of the H-bond energies on the helix length, as seen in Figure 4 and Table S1. The farther helical dipoles do not seem to affect the electronic structures of the H-bond donors and acceptors, although they contribute to the cooperative nature of the α-helix formation^23)^.

Thus, we can conclude that the H-bond energies of the α-helix, as in the AH and ST models, are generally weaker than those of the separated H-bonds, as in the MH model, due to the depolarized electronic structures around the carbonyl oxygen of the *i*-th residue and the amide hydrogen of the (*i*+4)-th residue. Such depolarizations redistribute the electron density, and are caused by the local electronic interactions in their neighborhood inside the α-helical structure. Similar H-bond energy changes depending on the peptide backbone structures were also found in the antiparallel β-sheet models in our previous paper^14)^ and others^27,28)^. When the SEs were computed, the odd-numbered β-sheet models had weaker SEs by forming smaller hydrogen bond ring structures, and the even-numbered β-sheet models had stronger SEs by forming larger hydrogen bond ring structures^14)^.

### Towards improvement of the H-bond energy by the classical force-field

The current analysis of the depolarization of the electronic structure at the carbonyl group of the *i*-th residue and the amide group of the (*i*+4)-th residue in an α-helix revealed the chemical origin of the incompleteness of any force-field parameters in MM computations for H-bond energies. Thus, we have the opportunity to overcome this problem by improving the force-field parameters to obtain more realistic H-bond energies.

In order to reproduce the actual H-bond energies by an MM computation, there are two putative ways to improve the force-fields, by modifying either the atomic partial charges or the backbone dihedral parameters.

The first approach that introduced new atomic partial charges, which are not constant but depend on the local molecular structures, is promising. Since the introduction of the classical MM computations and MD simulations, the constant atomic partial charges for backbone atoms have been widely used independently of the protein conformations, although the atomic charges greatly depend on the backbone structure. Numerous efforts have sought to develop force-fields, in which the polarization effects are included depending on the local electrostatic field^12, 13)^. However, there were no systematic approaches to develop the atomic partial charges of the backbone atoms, which depend on the local backbone structures of peptides and proteins.

In the second approach to create new backbone dihedral parameters, they should depend not only on a single backbone parameter set (*φ_i_, ψ_i_, ω_i_*) of the single *i*-th residue, but also on the (*φ*_*i*−1*1*_, *ψ*_*i*−1_, *ω*_*i*−1_) and (*φ*_*i*+*1*_, *ψ*_*i*+*1*_, *ω*_*i*+*1*_) values of the neighboring residues. So far, almost all of the dihedral parameters have been computed based on di-peptides, such as Ace-Ala-Nme, and thus they have ignored the changes in the electronic structures depending on the neighboring backbone structures.

Neither of the above approaches is simple, because the classical force-field artificially separates the whole peptide energy to individual energy terms, such as electrostatic energy and dihedral angle terms, which in principle strongly correlate with each other. Therefore, the balance between the many force-field parameters is essential, when trying to improve the H-bond energy in an α-helix by a classical MM computation, based on the depolarizing phenomenon found in this study.

## Conclusion

The H-bond interaction energies and the associated electron density changes in the α-helical structures were systematically analyzed by the MTA method^16)^, with high quality DFT and MM computations. MTA with the DFT computation is a powerful method to estimate the H-bond interaction energy in a large system, in which the H-bond donors and acceptors are linked to many other atoms with covalent bonds. The H-bond interaction energy in an α-helix depends strongly on the local backbone conformation, and the tendencies are well reproduced even by the classical MM model, based on the AMBER ff99SB force-field parameters^15)^.

We first prepared the α-helical peptide models (AH models) by energy minimized Ace-(Ala)_*n*_-Nme α-helical structures, where *n* ranged from 3 to 8. In order to quantitatively dissect the origin of the H-bond interaction energy of the α-helix, we constructed the minimal H-bond model (MH model), which is composed of only the atoms forming a single H-bond, and a single-turn model (ST model), which is composed of three successive alanine residues, (Ala)_3_, in the α-helix capped by acetyl and *N*-methyl groups at the N- and C-termini, respectively. The individual H-bond energies were computed by using MTA with the DFT method. We found that the H-bond energies of the AH and ST models were always significantly weaker than those of the MH model. Interestingly, the H-bond energy values of the MH model were similar to those of the MM model. The H-bond energies of the AH model were only slightly weaker than those of the ST model.

Our current Hirshfeld population analysis for the Ace-(Ala)_8_-Nme model structures suggested that due to the closely located electric dipole of the carbonyl (C=O) group at the (*i*+1)-th residue, *μ*^*i*+1^_CO_, to *μ^i^*_CO_ at the *i*-th residue in the parallel direction in an α-helix, the dipole moments should decrease by depolarization. Similarly, due to the closely located dipole of the amide (NH) group at the (*i*+3)-th residue, *μ*^*i*+3^_HN_, to *μ*^*i*+4^_HN_ at the (*i*+4)-th residue in the parallel direction, the dipole moments should also decrease. Thus, the local dipole moments, *μ^i^*_CO_ and *μ*^*i*+4^_HN_, were reduced by about 5−6% to 5−8%, respectively. Moreover, the contributions from another neighboring C=O group at the (*i*−1)-th residue, *μ*^*i*−1^_CO_, and the NH group at the (*i*+5)-th residue, *μ^i^^*, were also analyzed. Their local dipole moments were both reduced by about 2-3%.

So far, the MTA method has only been used to approximate the local energies in large systems^16)^. Here, we have shown that the electronic structures are also provided in the details of the MTA method, and that the distributions of electron densities and their changes upon H-bond formation in the α-helix are useful to reveal the chemical origins of the H-bond interaction energies.

Since the classical MM computations and MD simulations started to be employed many years ago, the constant atomic partial charges independent of the protein conformations have been widely used, although the atomic polarization effects were sometimes included depending on the local electrostatic field^12,13)^. However, the electronic structures can change depending on the local structures and the environments of peptides and proteins. Recent QM/MM and QM/MD simulations could overcome these issues at some local and focused areas^29–31)^, but they are not applicable to entire macromolecular systems, which include many α-helices. Thus, the H-bond energy values in MM computations should be improved by introducing new atomic partial charges or better backbone dihedral parameters, which should depend on the local peptide backbone structures. Newer and better force-fields than the current ones are required for more reliable molecular simulations, particularly for better understanding of intrinsically disordered regions in proteins, which are abundant and important in many biological systems^4,5,32,33)^.

## Acknowledgements

This work was supported by a MEXT Grant-in-Aid for Scientific Research on Innovative Areas “3D Active-Site Science” (26105012), by JSPS Grants-in-Aid for Exploratory Research (23657103 and JP16K14711), by a JSPS Grant-in-Aid for Scientific Research (C) (JP16K07325), and by JST CREST (JPMJCR14M3). The computations were performed at the Research Center for Computational Science, Okazaki, Japan. This work was performed in part under the Cooperative Research Program of the Institute for Protein Research, Osaka University, CR-15-05, CR-16-05, and CR-17-05. We thank Tomohiro Maruyama for the computations of the Hirshfeld charges.

## Supplementary Materials

**Table S1:** Hydrogen bond (H-bond) energies in α-helices for AH (α-helical structure) model, ST (single turn) model, and MH (minimal H-bond) model by the MTA method and those by the MM computation with AMBER ff99SB force field parameters. The stabilization energies, Δ*E*_SE_^total^, defined by eq. [4] are also listed. The energy minimized α-helix structures were constructed following the procedure mentioned in the Method section.

| α-helix Structure | Distance<sup>a)</sup><br>(Å) | CO-HN angle<sup>b)</sup><br>(degree) | H-bond Energy (kcal/mol) | | | | $\Delta E_{SE}^{total}$ h)<br>(kcal/mol) |
| --- | --- | --- | --- | --- | --- | --- | --- |
|  |  |  | AH model<sup>c)</sup> | ST model<sup>d)</sup> | MH model<sup>e)</sup> | MM<sup>f)</sup> |  |
| 3-1 | 2.418 | 171.796 | -2.875 | -2.870 | -4.018 | -4.078 | -4.880 |
| 4-1 | 2.363 | 175.977 | -2.980 | -3.296 | -4.509 | -4.425 | -5.361 |
| 4-2 | 2.343 | 167.386 | -2.786 | -2.969 | -4.004 | -3.839 | -4.925 |
| 5-1 | 2.332 | 178.102 | -3.352 | -3.519 | -4.780 | -4.622 | -5.639 |
| 5-2 | 2.268 | 168.123 | -2.911 | -3.420 | -4.533 | -4.296 | -5.458 |
| 5-3 | 2.421 | 164.590 | -2.470 | -2.567 | -3.492 | -3.257 | -4.471 |
| 6-1 | 2.313 | 178.503 | -3.652 | -3.549 | -4.829 | -4.707 | -5.695 |
| 6-2 | 2.199 | 168.542 | -3.362 | -3.713 | -4.891 | -4.631 | -5.827 |
| 6-3 | 2.345 | 163.525 | -2.693 | -3.087 | -4.078 | -3.744 | -5.046 |
| 6-4 | 2.366 | 167.549 | -2.856 | -2.806 | -3.832 | -3.672 | -4.788 |
| 7-1 | 2.292 | 178.062 | -3.765 | -3.585 | -4.880 | -4.792 | -5.752 |
| 7-2 | 2.172 | 168.146 | -3.657 | -3.748 | -4.955 | -4.752 | -5.895 |
| 7-3 | 2.292 | 163.714 | -3.126 | -3.378 | -4.408 | -4.009 | -5.385 |
| 7-4 | 2.293 | 167.155 | -3.020 | -3.291 | -4.394 | -4.154 | -5.336 |
| 7-5 | 2.345 | 168.055 | -3.001 | -2.867 | -3.902 | -3.753 | -4.862 |
| 8-1 | 2.273 | 177.763 | -3.902 | -3.632 | -4.945 | -4.878 | -5.821 |
| 8-2 | 2.147 | 168.060 | -3.789 | -3.781 | -5.004 | -4.851 | -5.948 |
| 8-3 | 2.256 | 163.435 | -3.422 | -3.431 | -4.484 | -4.157 | -5.473 |
| 8-4 | 2.220 | 167.585 | -3.504 | -3.588 | -4.753 | -4.495 | -5.708 |
| 8-5 | 2.263 | 167.404 | -3.187 | -3.350 | -4.472 | -4.264 | -5.422 |
| 8-6 | 2.339 | 167.613 | -3.031 | -2.867 | -3.895 | -3.731 | -4.866 |

a) Distance between O atom of C=O at *i*-th residue and H atom of HN at (*i*+4)-th residue
b) Angle formed by the two vectors of C=O at *i*-th residue and HN at (*i*+4)-th residue
c) H-bond energies by AH (original α-helical structure) model with MTA method
d) H-bond energies by ST (single turn) model with MTA method
e) H-bond energies by MH (minimal H-bond) model with MTA method
f) Computed by the MM method with AMBER ff99SB force field parameters
g) Computed by the MM method with the modified atomic charges for C=O at *i*-th residue and HN at (*i*+4)-th residue (See text and Table S3).
h) Stabilization energy defined by eq. [4]

**Table S2:**
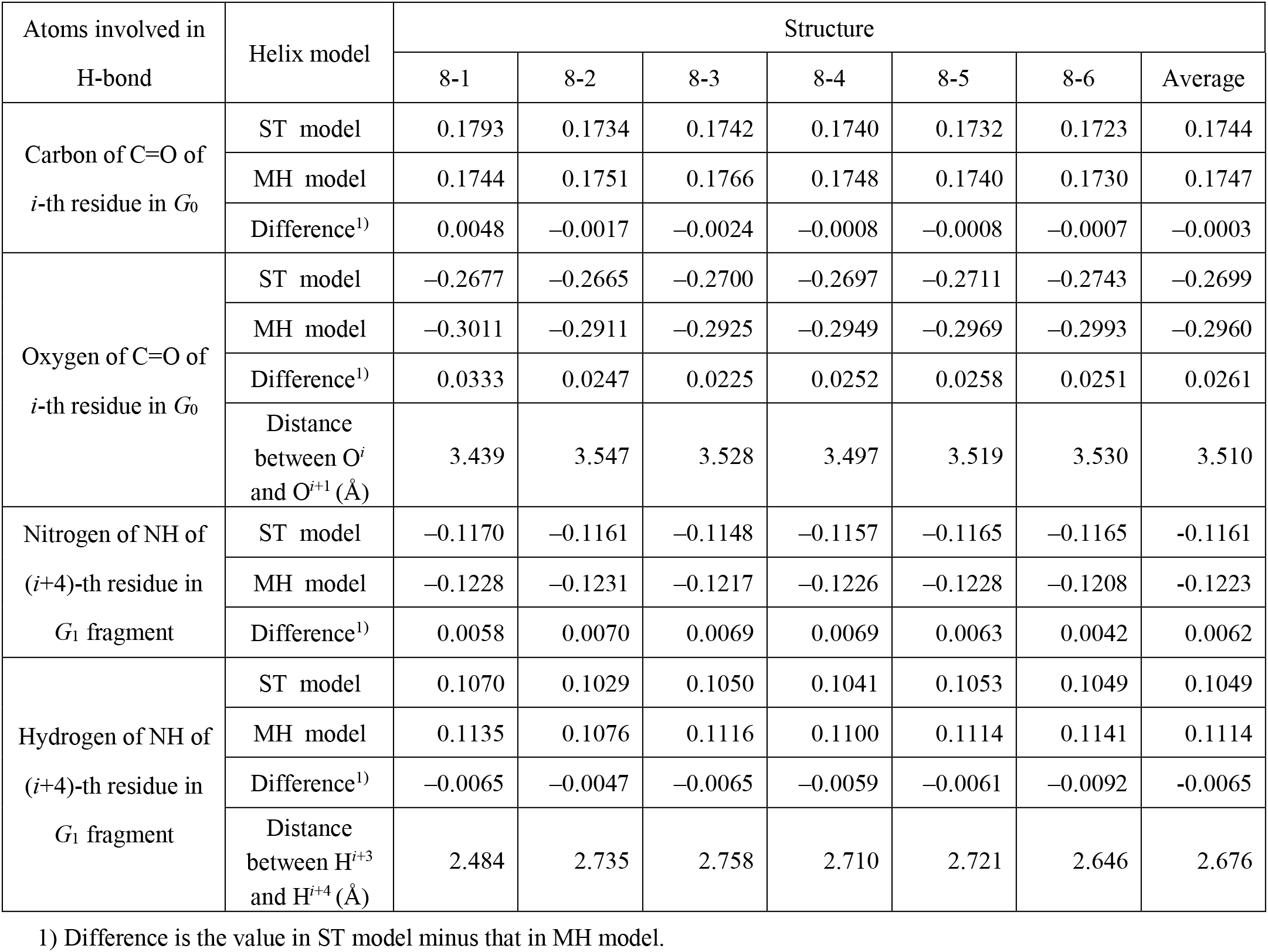
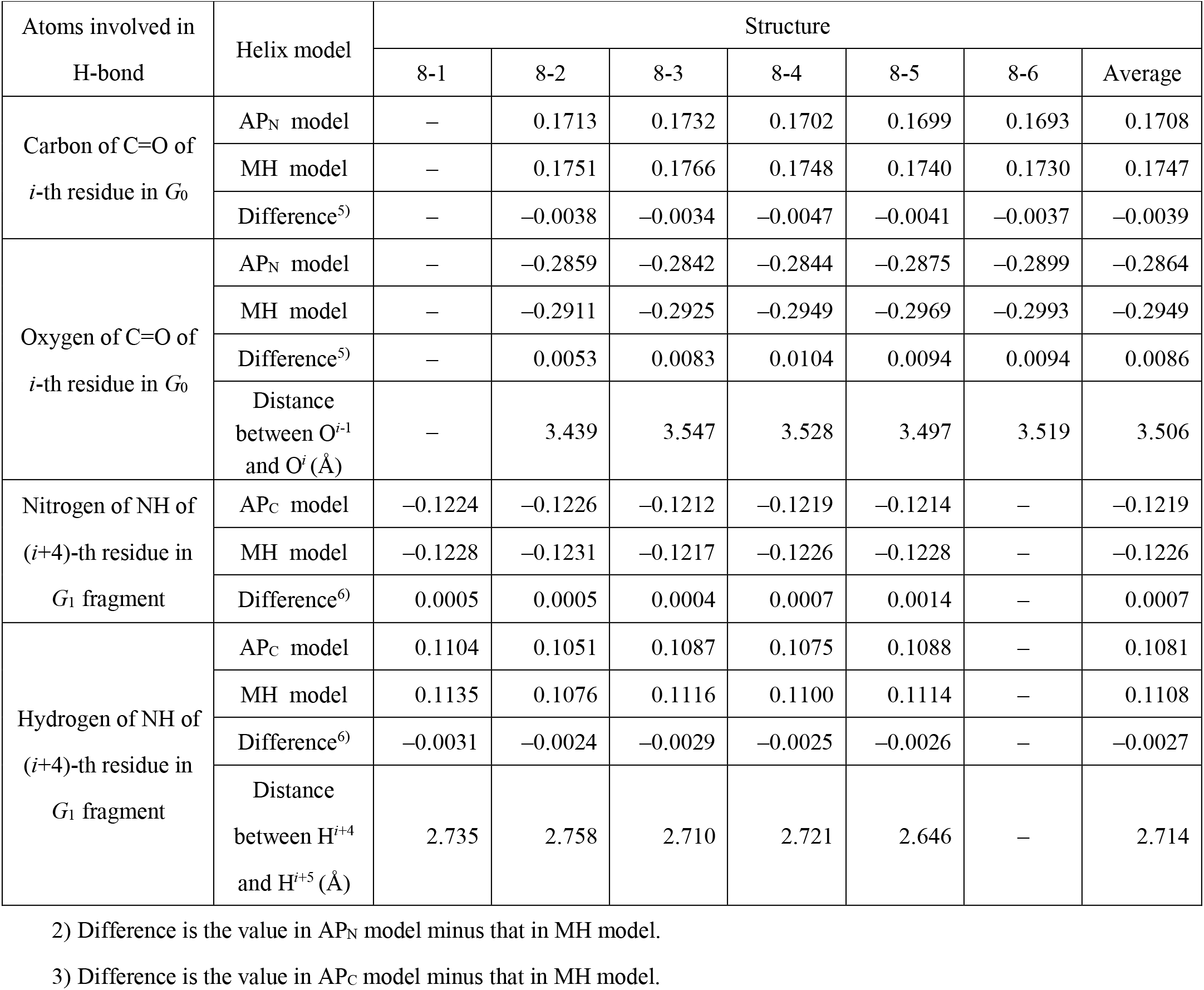
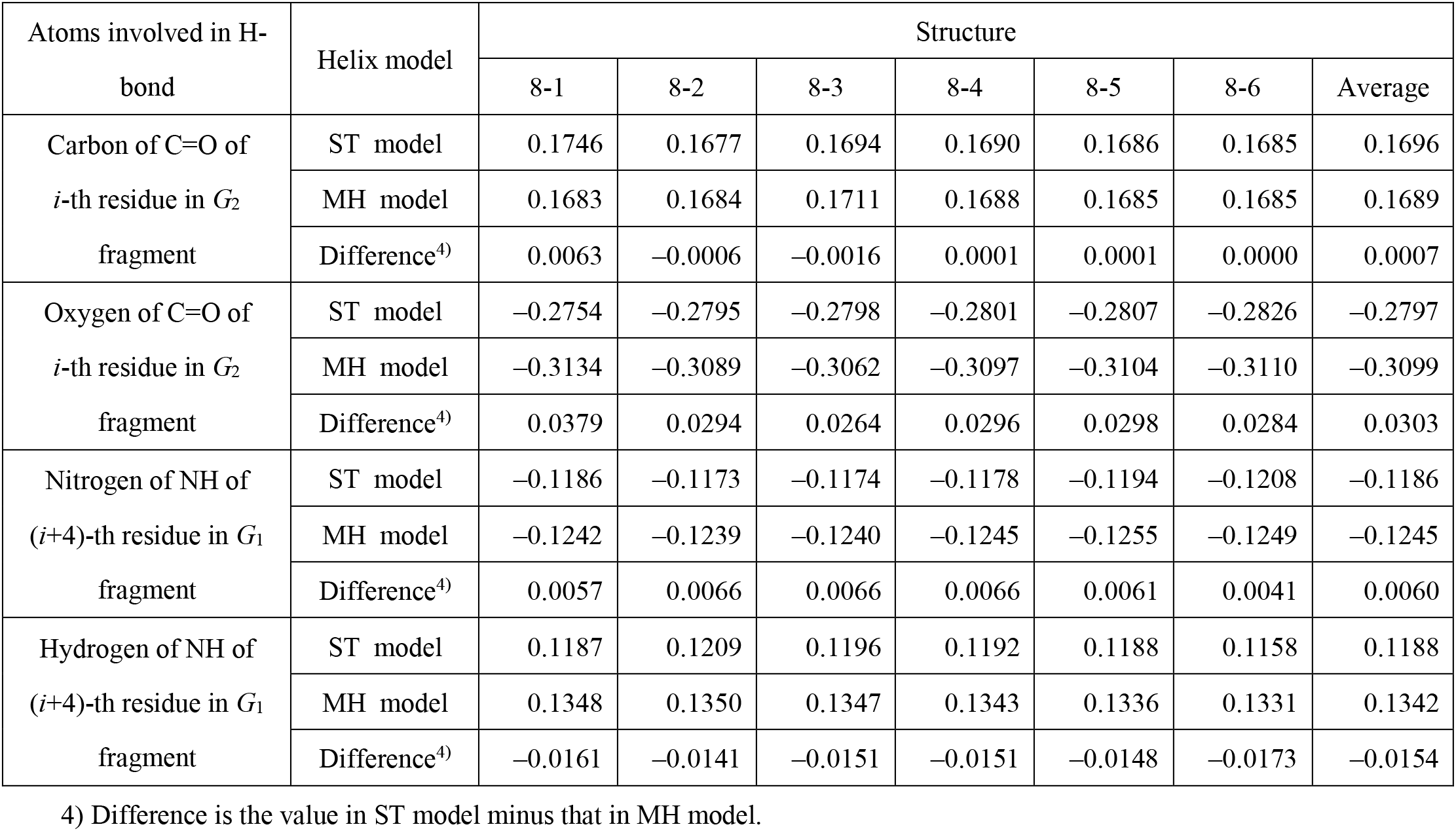
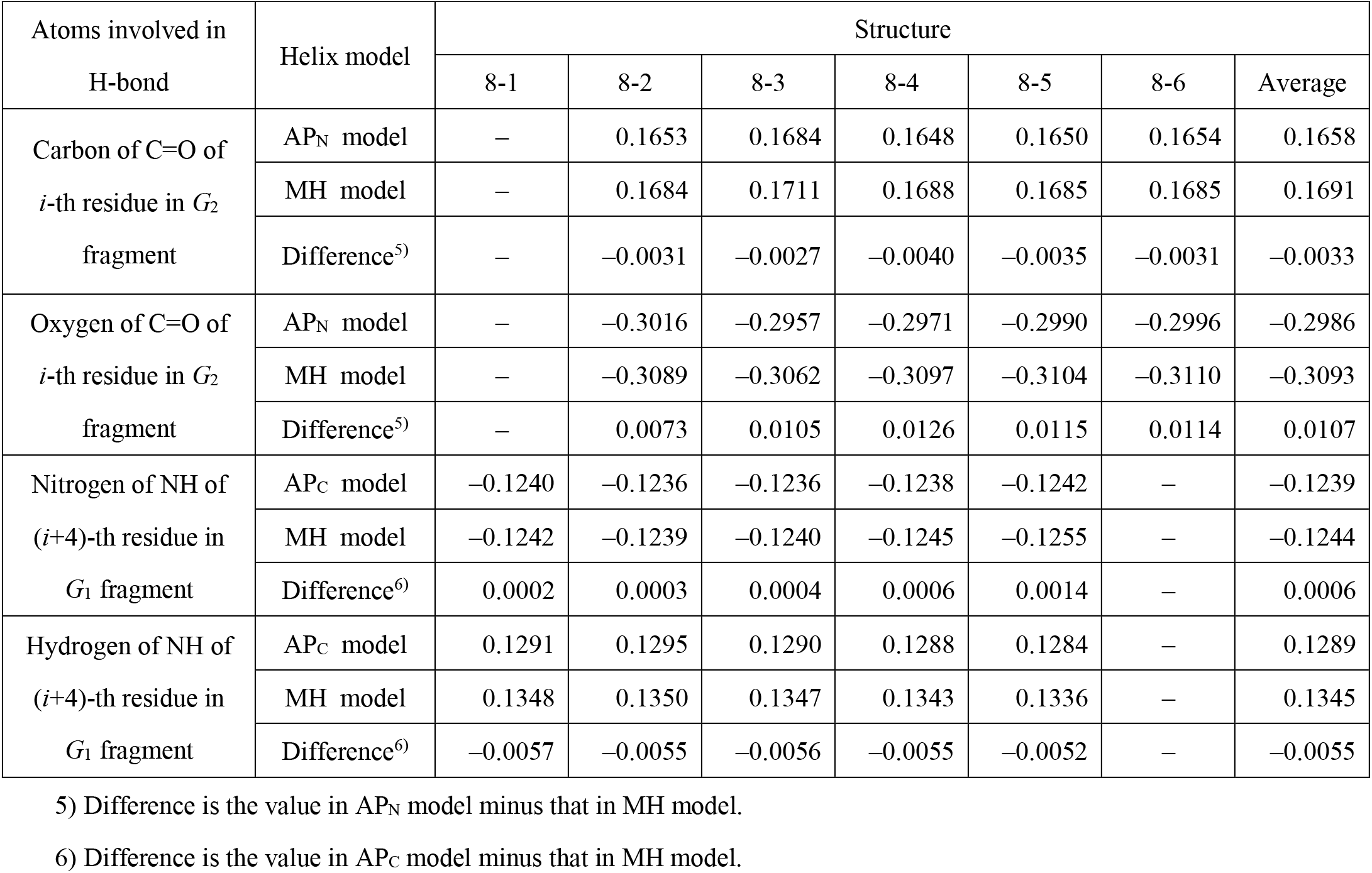
Hirshfeld atomic charges of the carbonyl group (C=O) of *i*-th residue and those of the amide group (NH) of (*i*+4)-th residue in ST, AP_N_, AP_C_ and MH models, for *G*_0_, *G*_1_ and *G*_2_ fragments (see Figure 2), respectively. The differences are also shown. The optimized six α-helix structures were constructed following the procedure mentioned in the Method section. A) Hirshfeld charges in *G*_0_, where H-bonds are formed between C=O of *i*-th residue and NH of (*i*+4)-th residue for ST and MH models B) Hirshfeld charges in *G*_0_, where H-bonds are formed between C=O of *i*-th residue and NH of (*i*+4)-th residue for AP_N_ and AP_C_ models C) Hirshfeld charges in *G*_1_ and *G*_2_ fragments, where no H-bonds are formed between C=O of *i*-th residue and NH of (*i*+4)-th residue for ST and MH models D) Hirshfeld charges in in *G*_1_ and *G*_2_ fragments, where no H-bonds are formed between C=O of *i*-th residue and NH of (*i*+4)-th residue for AP_N_ and AP_C_ models

| Atoms involved in H-bond | Helix model | Structure |  |  |  |  |  |  |
| --- | --- | --- | --- | --- | --- | --- | --- | --- |
|  |  | 8-1 | 8-2 | 8-3 | 8-4 | 8-5 | 8-6 | Average |
| Carbon of C=O of $i$-th residue in $G_0$ | ST model | 0.1793 | 0.1734 | 0.1742 | 0.1740 | 0.1732 | 0.1723 | 0.1744 |
|  | MH model | 0.1744 | 0.1751 | 0.1766 | 0.1748 | 0.1740 | 0.1730 | 0.1747 |
|  | Difference<sup>1)</sup> | 0.0048 | -0.0017 | -0.0024 | -0.0008 | -0.0008 | -0.0007 | -0.0003 |
| Oxygen of C=O of $i$-th residue in $G_0$ | ST model | -0.2677 | -0.2665 | -0.2700 | -0.2697 | -0.2711 | -0.2743 | -0.2699 |
|  | MH model | -0.3011 | -0.2911 | -0.2925 | -0.2949 | -0.2969 | -0.2993 | -0.2960 |
|  | Difference<sup>1)</sup> | 0.0333 | 0.0247 | 0.0225 | 0.0252 | 0.0258 | 0.0251 | 0.0261 |
|  | Distance between O<sup>*i*</sup> and O<sup>*i+1*</sup> (Å) | 3.439 | 3.547 | 3.528 | 3.497 | 3.519 | 3.530 | 3.510 |
| Nitrogen of NH of $(i+4)$-th residue in $G_1$ fragment | ST model | -0.1170 | -0.1161 | -0.1148 | -0.1157 | -0.1165 | -0.1165 | -0.1161 |
|  | MH model | -0.1228 | -0.1231 | -0.1217 | -0.1226 | -0.1228 | -0.1208 | -0.1223 |
|  | Difference<sup>1)</sup> | 0.0058 | 0.0070 | 0.0069 | 0.0069 | 0.0063 | 0.0042 | 0.0062 |
| Hydrogen of NH of $(i+4)$-th residue in $G_1$ fragment | ST model | 0.1070 | 0.1029 | 0.1050 | 0.1041 | 0.1053 | 0.1049 | 0.1049 |
|  | MH model | 0.1135 | 0.1076 | 0.1116 | 0.1100 | 0.1114 | 0.1141 | 0.1114 |
|  | Difference<sup>1)</sup> | -0.0065 | -0.0047 | -0.0065 | -0.0059 | -0.0061 | -0.0092 | -0.0065 |
|  | Distance between H<sup>*i+3*</sup> and H<sup>*i+4*</sup> (Å) | 2.484 | 2.735 | 2.758 | 2.710 | 2.721 | 2.646 | 2.676 |

1) Difference is the value in ST model minus that in MH model.

| Atoms involved in H-bond | Helix model | Structure |  |  |  |  |  |  |
| --- | --- | --- | --- | --- | --- | --- | --- | --- |
|  |  | 8-1 | 8-2 | 8-3 | 8-4 | 8-5 | 8-6 | Average |
| Carbon of C=O of $i$-th residue in $G_0$ | AP<sub>N</sub> model | – | 0.1713 | 0.1732 | 0.1702 | 0.1699 | 0.1693 | 0.1708 |
|  | MH model | – | 0.1751 | 0.1766 | 0.1748 | 0.1740 | 0.1730 | 0.1747 |
|  | Difference<sup>5)</sup> | – | –0.0038 | –0.0034 | –0.0047 | –0.0041 | –0.0037 | –0.0039 |
| Oxygen of C=O of $i$-th residue in $G_0$ | AP<sub>N</sub> model | – | –0.2859 | –0.2842 | –0.2844 | –0.2875 | –0.2899 | –0.2864 |
|  | MH model | – | –0.2911 | –0.2925 | –0.2949 | –0.2969 | –0.2993 | –0.2949 |
|  | Difference<sup>5)</sup> | – | 0.0053 | 0.0083 | 0.0104 | 0.0094 | 0.0094 | 0.0086 |
|  | Distance between O<sup>$i-1$</sup> and O<sup>$i$</sup> (Å) | – | 3.439 | 3.547 | 3.528 | 3.497 | 3.519 | 3.506 |
| Nitrogen of NH of $(i+4)$-th residue in $G_1$ fragment | AP<sub>C</sub> model | –0.1224 | –0.1226 | –0.1212 | –0.1219 | –0.1214 | – | –0.1219 |
|  | MH model | –0.1228 | –0.1231 | –0.1217 | –0.1226 | –0.1228 | – | –0.1226 |
|  | Difference<sup>6)</sup> | 0.0005 | 0.0005 | 0.0004 | 0.0007 | 0.0014 | – | 0.0007 |
| Hydrogen of NH of $(i+4)$-th residue in $G_1$ fragment | AP<sub>C</sub> model | 0.1104 | 0.1051 | 0.1087 | 0.1075 | 0.1088 | – | 0.1081 |
|  | MH model | 0.1135 | 0.1076 | 0.1116 | 0.1100 | 0.1114 | – | 0.1108 |
|  | Difference<sup>6)</sup> | –0.0031 | –0.0024 | –0.0029 | –0.0025 | –0.0026 | – | –0.0027 |
|  | Distance between H<sup>$i+4$</sup> and H<sup>$i+5$</sup> (Å) | 2.735 | 2.758 | 2.710 | 2.721 | 2.646 | – | 2.714 |
|  |  | 8-1 | 8-2 | 8-3 | 8-4 | 8-5 | 8-6 | Average |
| Carbon of C=O of $i$-th residue in $G_2$ fragment | ST model | 0.1746 | 0.1677 | 0.1694 | 0.1690 | 0.1686 | 0.1685 | 0.1696 |
|  | MH model | 0.1683 | 0.1684 | 0.1711 | 0.1688 | 0.1685 | 0.1685 | 0.1689 |
|  | Difference<sup>4)</sup> | 0.0063 | -0.0006 | -0.0016 | 0.0001 | 0.0001 | 0.0000 | 0.0007 |
| Oxygen of C=O of $i$-th residue in $G_2$ fragment | ST model | -0.2754 | -0.2795 | -0.2798 | -0.2801 | -0.2807 | -0.2826 | -0.2797 |
|  | MH model | -0.3134 | -0.3089 | -0.3062 | -0.3097 | -0.3104 | -0.3110 | -0.3099 |
|  | Difference<sup>4)</sup> | 0.0379 | 0.0294 | 0.0264 | 0.0296 | 0.0298 | 0.0284 | 0.0303 |
| Nitrogen of NH of $(i+4)$-th residue in $G_1$ fragment | ST model | -0.1186 | -0.1173 | -0.1174 | -0.1178 | -0.1194 | -0.1208 | -0.1186 |
|  | MH model | -0.1242 | -0.1239 | -0.1240 | -0.1245 | -0.1255 | -0.1249 | -0.1245 |
|  | Difference<sup>4)</sup> | 0.0057 | 0.0066 | 0.0066 | 0.0066 | 0.0061 | 0.0041 | 0.0060 |
| Hydrogen of NH of $(i+4)$-th residue in $G_1$ fragment | ST model | 0.1187 | 0.1209 | 0.1196 | 0.1192 | 0.1188 | 0.1158 | 0.1188 |
|  | MH model | 0.1348 | 0.1350 | 0.1347 | 0.1343 | 0.1336 | 0.1331 | 0.1342 |
|  | Difference<sup>4)</sup> | -0.0161 | -0.0141 | -0.0151 | -0.0151 | -0.0148 | -0.0173 | -0.0154 |

4) Difference is the value in ST model minus that in MH model.

| Atoms involved in H-bond | Helix model | Structure |  |  |  |  |  |  |
| --- | --- | --- | --- | --- | --- | --- | --- | --- |
|  |  | 8-1 | 8-2 | 8-3 | 8-4 | 8-5 | 8-6 | Average |
| Carbon of C=O of $i$-th residue in $G_2$ fragment | AP<sub>N</sub> model | – | 0.1653 | 0.1684 | 0.1648 | 0.1650 | 0.1654 | 0.1658 |
|  | MH model | – | 0.1684 | 0.1711 | 0.1688 | 0.1685 | 0.1685 | 0.1691 |
|  | Difference<sup>5)</sup> | – | –0.0031 | –0.0027 | –0.0040 | –0.0035 | –0.0031 | –0.0033 |
| Oxygen of C=O of $i$-th residue in $G_2$ fragment | AP<sub>N</sub> model | – | –0.3016 | –0.2957 | –0.2971 | –0.2990 | –0.2996 | –0.2986 |
|  | MH model | – | –0.3089 | –0.3062 | –0.3097 | –0.3104 | –0.3110 | –0.3093 |
|  | Difference<sup>5)</sup> | – | 0.0073 | 0.0105 | 0.0126 | 0.0115 | 0.0114 | 0.0107 |
| Nitrogen of NH of $(i+4)$-th residue in $G_1$ fragment | AP<sub>C</sub> model | –0.1240 | –0.1236 | –0.1236 | –0.1238 | –0.1242 | – | –0.1239 |
|  | MH model | –0.1242 | –0.1239 | –0.1240 | –0.1245 | –0.1255 | – | –0.1244 |
|  | Difference<sup>6)</sup> | 0.0002 | 0.0003 | 0.0004 | 0.0006 | 0.0014 | – | 0.0006 |
| Hydrogen of NH of $(i+4)$-th residue in $G_1$ fragment | AP<sub>C</sub> model | 0.1291 | 0.1295 | 0.1290 | 0.1288 | 0.1284 | – | 0.1289 |
|  | MH model | 0.1348 | 0.1350 | 0.1347 | 0.1343 | 0.1336 | – | 0.1345 |
|  | Difference<sup>6)</sup> | –0.0057 | –0.0055 | –0.0056 | –0.0055 | –0.0052 | – | –0.0055 |

5) Difference is the value in AP<sub>N</sub> model minus that in MH model.
6) Difference is the value in AP<sub>C</sub> model minus that in MH model.

**Table S3:** H-bond energies in α-helices for HT_N_, HT_C_, AP_N_, and AP_C_ models computed by the MTA method with DFT.

| α-helix<br>Structure | H-bond Energy (kcal/mol) | | | |
| --- | --- | --- | --- | --- |
|  | HT<sub>N</sub><br>model<sup>a)</sup> | HT<sub>C</sub><br>model<sup>b)</sup> | AP<sub>N</sub><br>model<sup>c)</sup> | AP<sub>C</sub><br>model<sup>d)</sup> |
| 8-1 | -4.418 | -3.993 | – | -4.516 |
| 8-2 | -4.526 | -4.152 | -4.793 | -4.495 |
| 8-3 | -4.044 | -3.785 | -4.188 | -4.006 |
| 8-4 | -4.273 | -3.959 | -4.408 | -4.273 |
| 8-5 | -4.001 | -3.714 | -4.165 | -4.043 |
| 8-6 | -3.432 | -3.227 | -3.610 | - |

a) N-terminal Half-Turn model shown in Figure S5 (A).
b) HT<sub>C</sub> (C-terminal Half-Turn) model shown in Figure S5 (B).
c) AP<sub>N</sub> (Additional N-terminal Peptide) model shown in Figure S5 (C). Because the AP<sub>N</sub> model was built from Ace-(Ala)<sub>8</sub>-Nme, there was no model for 8-1.
d) AP<sub>C</sub> (Additional C-terminal Peptide) model shown in Figure S5 (D). . Because the AP<sub>C</sub> model was built from Ace-(Ala)<sub>8</sub>-Nme, there was no model for 8-6.

**Figure S1:**
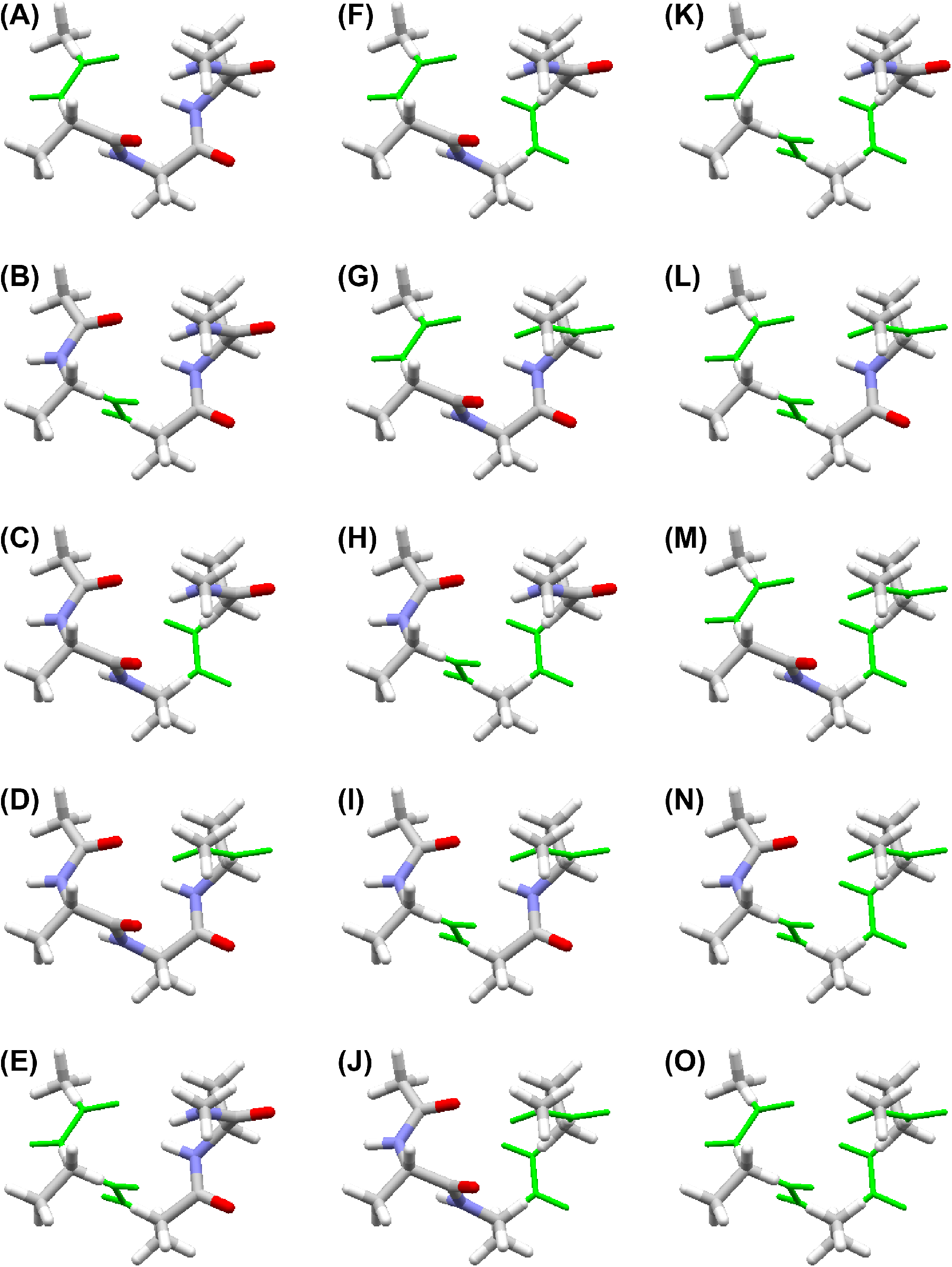
Fragment structures and their energies shown in the parentheses (kcal/mol) used in the current MTA method for Ace-(Ala)_3_-Nme: (A) model *F*_1_ (−822.4756), (B) *F*_2_ (−822.4808), (C) *F*_3_ (−822.4831), (D) *F*_4_ (−822.4831), (E) *F*_12_ (−655.0602), (F) *F*_13_ (−655.0649), (G) *F*_14_ (−655.0679), (H) *F*_23_ (−655.0669), (I) *F*_24_ (−655.0714), (J) *F*_34_ (−655.0670), (K) *F*_123_ (−487.6508), (L) *F*_124_ (−487.6559), (M) *F*_134_ (−487.6568), (N) *F*_234_ (−487.6562), (O) *F*_1234_ (−320.2462). Those energies were used in eq. [1] to compute the total MTA energy. The thin green lines are the original Ace-(Ala)_3_-Nme.

**Figure S2:**
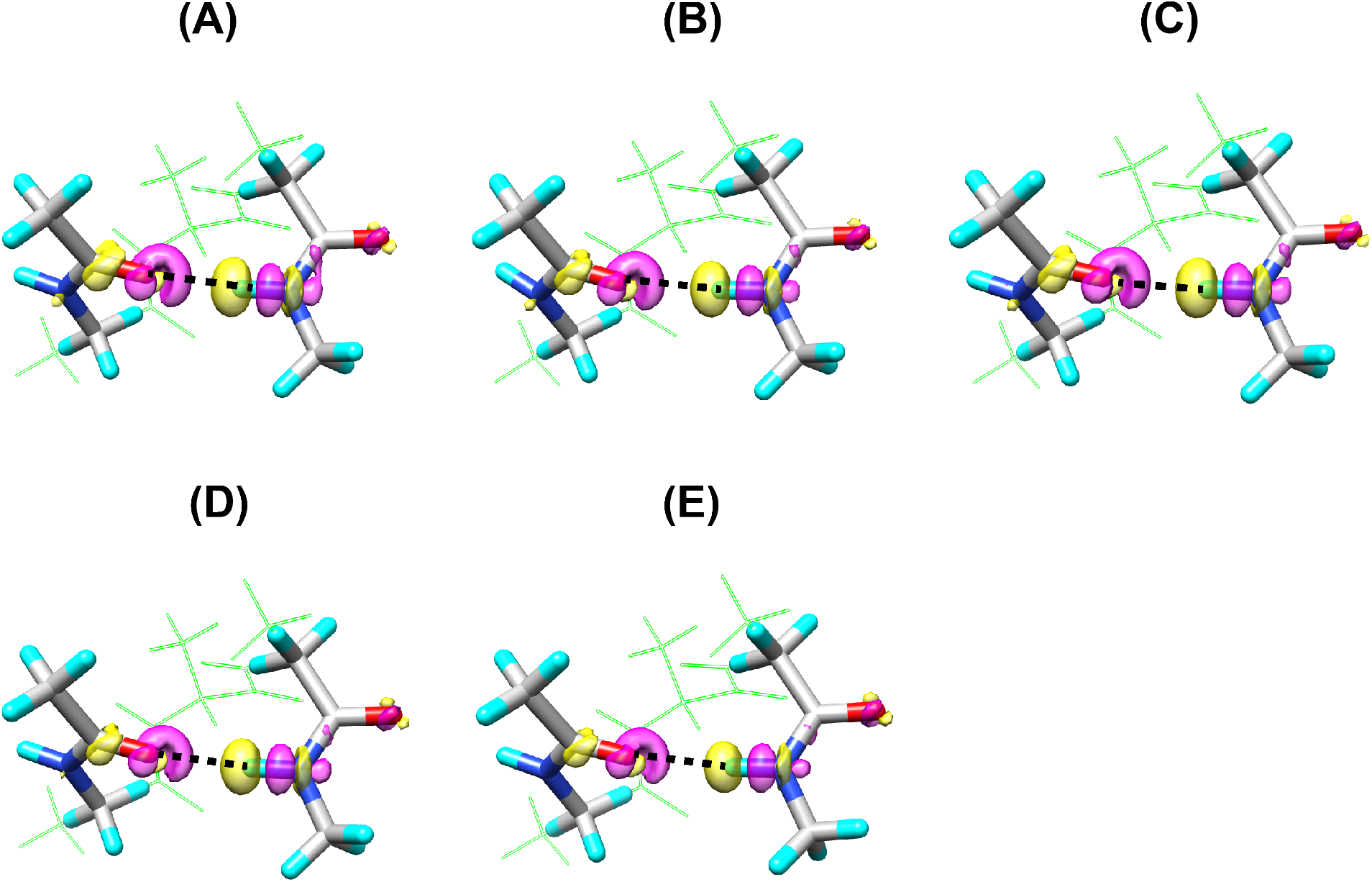
Electron density change upon H-bond formation Δ*ρ*_MTA_ given by eq. [3] in AH models for structures of (A) 8-2, (B) 8-3, (C) 8-4, (D) 8-5, and (E) 8-6. Yellow surface is the contour surface at −0.001 au, and magenta one is that at 0.001 au. The atoms in MH models are shown by stick model and the other atoms in ST models are shown by open green sticks. Each black dotted line is the H-bond between the oxygen atom of C=O group at *i*-th residue and the hydrogen atom of N-H group at (*i*+4)-th residue.

**Figure S3:**
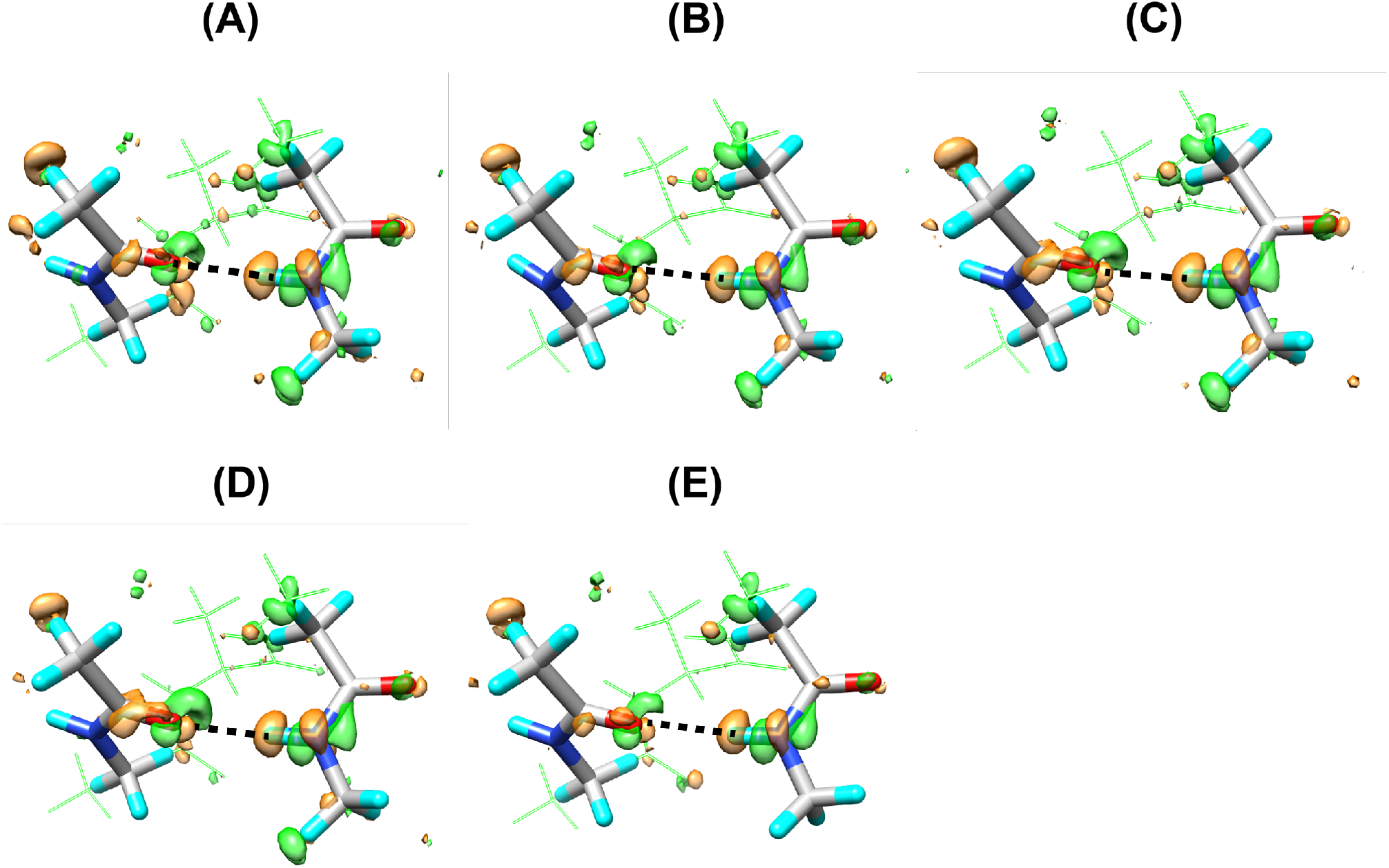
Difference in the electron density change upon H-bond formation between AH and MH models ΔΔ*ρ*_MTA_^AH−MH^ by eq. [7] for structures of (A) 8-2, (B) 8-3, (C) 8-4, (D) 8-5, and (E) 8-6. Green surfaces are the contour surfaces at −0.0002 au, and orange ones are those at 0.0002 au. The atoms in MH models are shown by sticks with CPK colors, and other atoms in ST models are by open green sticks. The black dotted lines are the H-bonds between the oxygen atoms of C=O groups at *i*-th residues and the hydrogen atoms of N-H groups at (*i*+4)-th residues.

**Figure S4:**
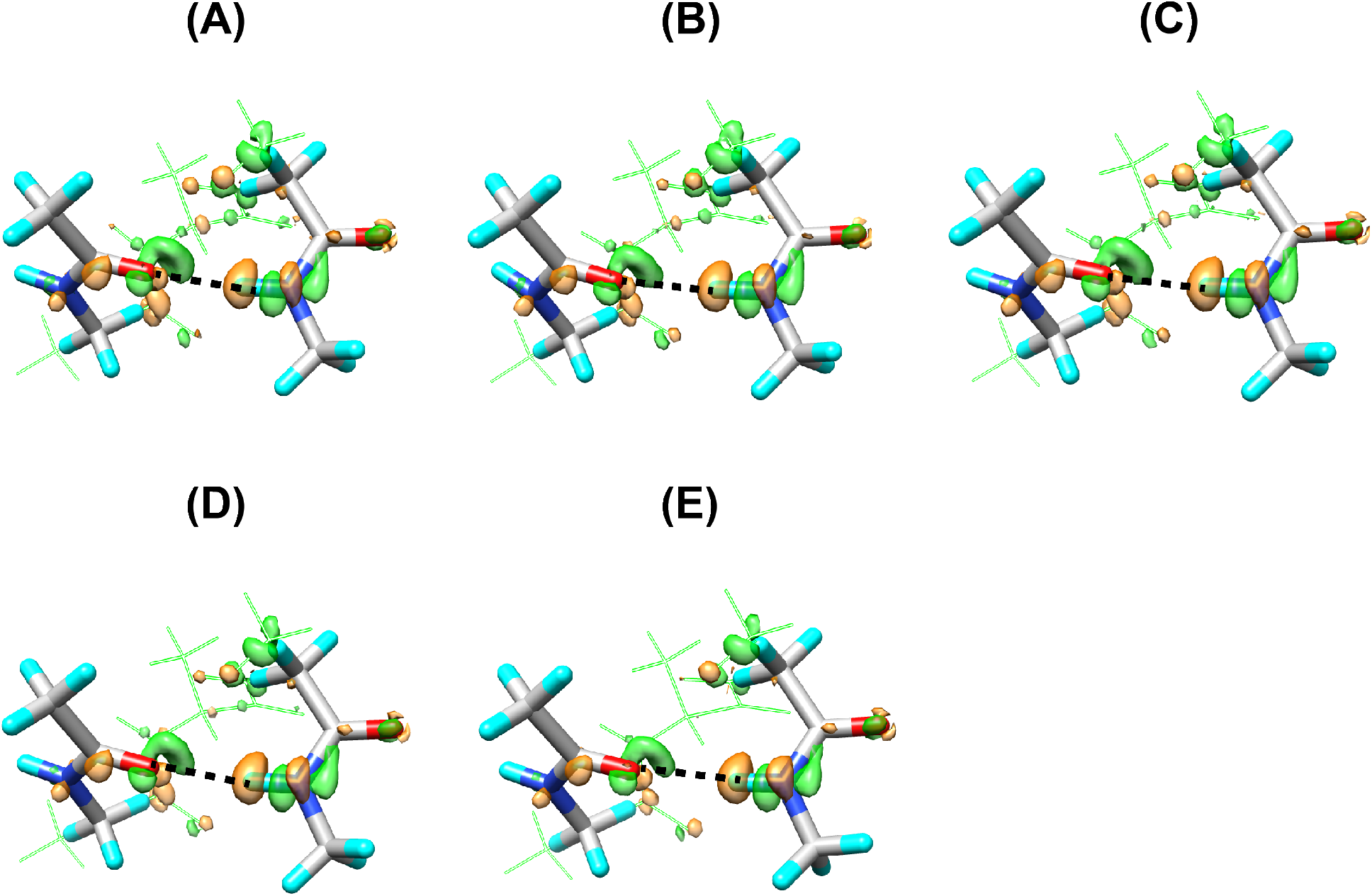
Difference in the electron density change upon H-bond formation between ST and MH models ΔΔ*ρ*_MTA_^ST−MH^ by eq. [7] for structures of (A) 8-2, (B) 8-3, (C) 8-4, (D) 8-5, and (E) 8-6. Green surfaces are the contour surfaces at −0.0002 au, and orange ones are those at 0.0002 au. The atoms in MH models are shown by sticks with CPK colors, and other atoms in ST models are by open green sticks. The black dotted lines are the H-bonds between the oxygen atoms of C=O groups at *i*-th residues and the hydrogen atoms of N-H groups at (*i*+4)-th residues.

**Figure S5:**
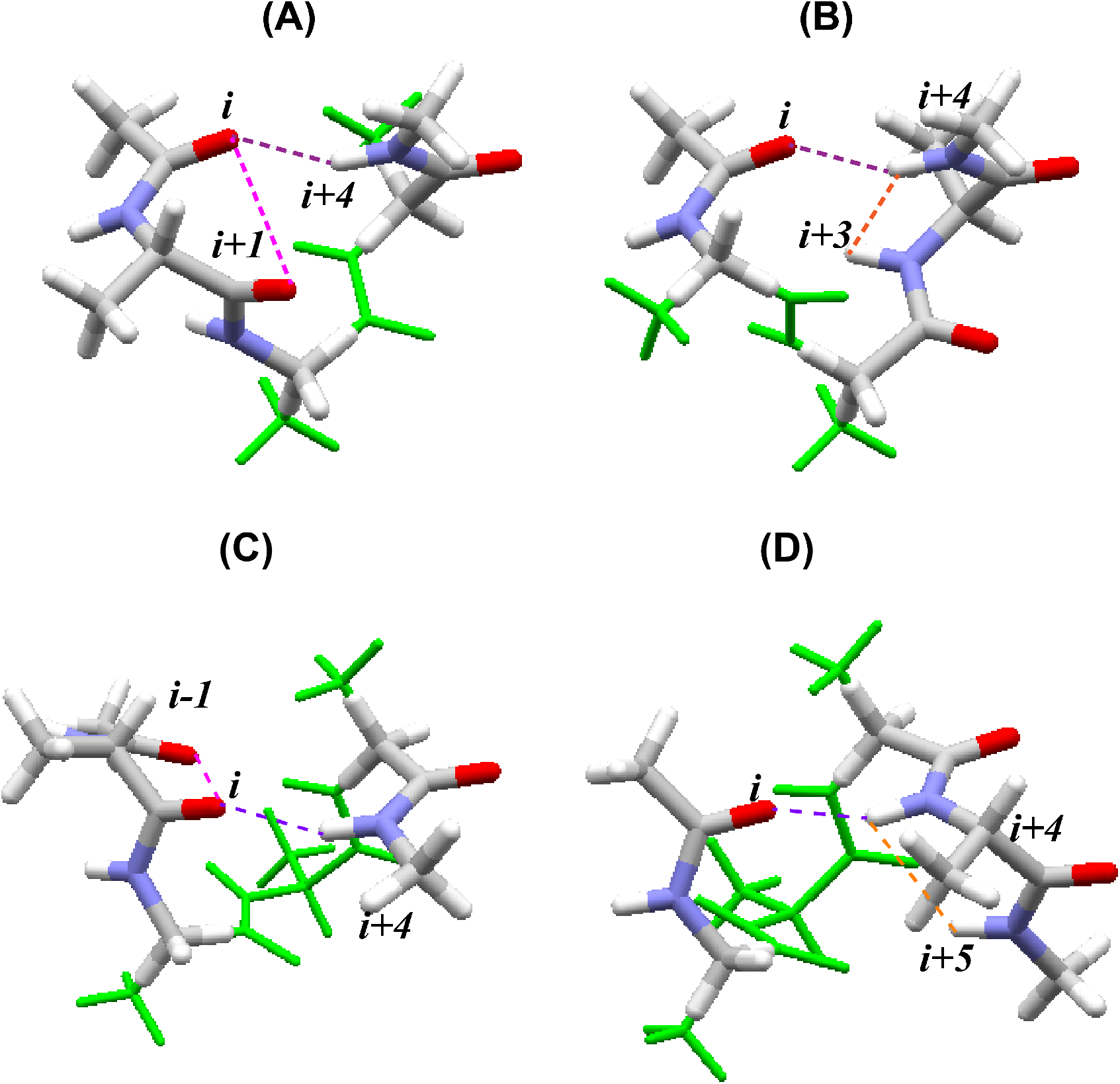
(A) HT_N_ (N-terminal Half-Turn) model, and (B) HT_C_ (C-terminal Half-Turn) model. The N-terminus of HT_N_ model and the C-terminus of HT_C_ model are both Ace-Ala-Nme, instead of the minimal Ace-Nme groups. Thus, HT_N_ model has an interaction between the successive C=O groups at *i*-th and (*i*+1)-th residues indicated by a magenta dotted line in (A), and HT_C_ model has another interaction between the successive NH groups at (*i*+3)-th and (*i*+4)-th residues indicated by an orange dotted line in (B). The target H-bonds between C=O groups at *i*-th residue and NH groups at (*i*+4)-th residues are indicated by purple dotted lines. The thin green lines are the original ST model, Ace- (Ala)_3_-Nme. (C) AP_N_ (N-terminal Additional-Peptide) model, and (D) AP_C_ (C-terminal Additional-Peptide) model. Ace-Ala group and Ala-Nme group are added at the N-terminus of AP_N_ model and the C-terminus of AP_C_ model, respectively. Thus, AP_N_ model has an interaction between the successive C=O groups at (*i*−1)-th and *i*-th residues indicated by a magenta dotted line in (C), and AP_C_ model has another interaction between the successive NH groups at (*i*+4)-th and (*i*+5)-th residues indicated by an orange dotted line in (D). The target H-bonds between C=O groups at *i*-th residue and NH groups at (*i*+4)-th residues are indicated by purple dotted lines. The thin green lines are the original ST model, Ace-(Ala)_3_-Nme.

